# Epidural Electrical Stimulation of the Cervical Dorsal Roots Restores Voluntary Upper Limb Control in Paralyzed Monkeys

**DOI:** 10.1101/2020.11.13.379750

**Authors:** B. Barra, S. Conti, M.G. Perich, K. Zhuang, G. Schiavone, F. Fallegger, K. Galan, N. D. James, Q. Barraud, M. Delacombaz, M. Kaeser, E. M. Rouiller, T. Milekovic, S. Lacour, J. Bloch, G. Courtine, M. Capogrosso

## Abstract

Recovering arm control is a top priority for people with paralysis. Unfortunately, the complexity of the neural mechanisms underlying arm control practically limited the effectiveness of neurotechnology approaches. Here, we exploited the neural function of surviving spinal circuits to restore voluntary arm and hand control in three monkeys with spinal cord injury using spinal cord stimulation. Our neural interface leverages the functional organization of the dorsal roots to convey artificial excitation via electrical stimulation to relevant spinal segments at appropriate movement phases. Stimulation bursts targeting specific spinal segments produced sustained arm movements enabling monkeys with arm paralysis to perform an unconstrained reach-and-grasp task. Stimulation specifically improved strength, task performances and movement quality. Electrophysiology suggested that residual descending inputs were necessary to produce coordinated movements. The efficacy and reliability of our approach hold realistic promises of clinical translation.

## INTRODUCTION

More than 5 million people in the US currently live with some form of motor paralysis^1^. Stroke and spinal cord injury (SCI) are the main causes with hundreds of thousands of new cases per year^2^. Impairments of the hand and arm are particularly problematic, representing a major unmet need for both SCI and stroke patient populations^3,4^. Indeed, even mild deficits in hand function lead to significant degradation of quality of life. Unfortunately, recovery of hand and arm motor function is still an unsolved clinical challenge.

Generated in the cerebral cortex, upper limb motor commands are relayed to subcortical and spinal circuits that activate motoneurons and regulate sensory inputs to produce skilled motor actions^5–8^. Spinal cord injury (SCI), or stroke, damage these communication pathways generating impairments in sensory regulation and motor functions that lead to motor paralysis.

Historically, neurotechnologies were conceived around the idea of restoring movements in paralyzed subjects via a technological bypass. Such solution would use signals from cortical areas as inputs and artificially compensate for lack of motoneuron activation by producing desired muscle activity below the lesion^9^. For example, functional electrical stimulation (FES) was used to activate arm muscles in response to intracortical neural activity from the motor cortex^10,11^. This pioneering concept allowed paralyzed monkeys and humans to perform voluntary grasping tasks^10–13^. However, translation of these concepts into daily clinical practice is hindered by two distinct limitations. First, the artificial motoneuron recruitment order generated by FES induces muscle fatigue^14^ which is particularly problematic for arm movements. Indeed, fatigue prevents the generation of sustained forces and consequently FES fails to enable sustained three-dimensional arm movements that are required for daily activities. Second, since FES bypasses surviving circuits in the spinal cord, complex stimulation protocols^15^ and sophisticated decoding algorithms^10,13^ are required to orchestrate the activation of multiple muscles and produce functional movements. As a result, these systems require an articulated combination of hardware and software. Unfortunately, this complexity does not cope well with dynamic clinical environments that need robust and practical solutions for a rapid set up and large-scale use.

In contrast, epidural electrical stimulation (EES) of the lumbar spinal cord exploits surviving spinal circuits and supra-spinal connections after injury to produce movements^16^. Similar to intraspinal stimulation^17–19^, EES engages motoneurons via direct recruitment of large sensory afferents^20,21^ leading to widespread excitatory post-synaptic potentials in the spinal cord. More importantly, since motoneurons are recruited via natural synaptic inputs, EES generates a natural recruitment order^22,23^ that is resistant to artificial fatigue. This enables the production of forces that can sustain the whole-body weight^24^. Moreover, engagement of motoneurons from pre-synaptic pathways allows residual descending inputs and spinal circuits to control motoneurons excitability and produce voluntary movement after complete motor paralysis^25,26^.

Building on animal models^27–29^, recent clinical studies have shown that continuous stimulation delivered through epidural implants on the dorsal aspect of the lumbosacral spinal cord increased muscle strength, voluntary muscle activation and single joint movements in people with complete leg paralysis^26,30,31^. More strikingly, when coupled with targeted physical rehabilitation protocols, continuous EES restored weight bearing locomotion in subjects with severe SCI^32,33^. These outstanding clinical results prompted experimental studies aiming at verifying whether EES could be used to promote also upper limb movements after SCI^34^. Unfortunately, while clinical studies showed some success in improving hand grip force with both epidural and non-invasive approaches^35,36^, continuous EES did not produce results of similar outstanding efficacy as those observed for the lower limbs^32,33^. In fact, clinical outcomes were similar to those obtained with surface FES^37^.

Reasons for this discrepancy may stem from the complexity of upper limb motor control and biomechanics compared to locomotion. Indeed, in contrast to pattern-driven^38,39^ and repetitive locomotor movements, upper limb movements are composed by a non-repetitive and task-dependent combination of movement modules which are highly dependent from sophisticated cortico-spinal control^7,40–44^ and accurate sensory feedback^42,45–47^. Because of this intrinsic complexity, non-specific neuromodulation could limit the efficacy of EES by exciting all spinal segments simultaneously, irrespectively of movement phase. More importantly, unspecific and continuous stimulation of the sensory afferents through EES disrupts natural sensory inputs^23^ thus hindering spinal regulation of movements which is critical in dexterous upper limb control^45–47^.

We and others have shown that it is possible to direct electrical stimulation of the spinal cord to target restricted segments during appropriate times^17,48,49^. These spatio-temporal stimulation protocols enabled voluntary locomotion in monkeys with SCI as early as day 6 post injury without any physical training^50^ and within 2 weeks post implantation in humans with complete leg paralysis^51^. This approach exploits the somato-topography of the spinal sensory system to selectively engage restricted spinal regions^21,49^. Unfortunately, non-invasive technologies and clinically approved electrodes are unfit for this scope^52,53^ because of their limits in selectivity. Therefore, we hypothesized that a neural interface, specifically designed to target the cervical dorsal roots, could enable the administration of spatio-temporal stimulation patterns to the cervical spinal cord. We tested this hypothesis in three monkeys with a unilateral cervical SCI. We designed a personalized epidural interface to target primary afferents within the cervical dorsal roots. We hypothesized that the electrical stimulation of the roots with bursts linked to movement attempts would enable voluntary motor control and improve functional deficits of the arm and hand that emerge after SCI. Specifically we tested for improvements in muscle strength, dexterity and ability to execute three-dimensional functional tasks in full independence. Finally, we verified that the mechanisms enabling the voluntary recruitment of motoneurons in the cervical spinal cord were similar to those occurring during EES of the lumbosacral circuits.

## Results

### Natural arm movements

Clinically effective systems should enable truly functional arm movements rather than simplified tasks such as single-joint movements. A functional arm movement entails a coordinated activation of arm muscles to achieve a desired movement while supporting the arm weight at all times. Most of daily activities require arm extension (reach) and flexion (pull), combined with a hand-grasp without a constrained timing or structure. Consequently, we developed a robotic platform allowing the quantification of reach, grasp and pull movements^54^ that would feel natural and unconstrained to monkeys both in trajectory and timings (**Figure 1A**). We trained three adult Macaca fascicularis monkeys to reach for, grasp, and pull an instrumented object placed on the end effector of our robotic arm (**Figure 1B**). Movement trajectories were not constrained neither kinematically nor in time. Monkeys waited for the go signal, reached for the object and pulled to receive a food or juice reward when the object crossed a pre-defined displacement threshold^54^. Monkeys intuitively and rapidly^29,30^ learned this task by developing their own individual kinematic strategies (**Extended Data Figure 1**) and personal movement speeds. We then designed a battery of electrophysiology and kinematic measurements to evaluate functional outcomes on task performances, muscle activation, muscle strength and movement dexterity. Specifically, we quantified full-limb 3D kinematics (Vicon Motion Systems, Oxford, UK), pulling forces, and electromyographic (EMG) signals from intramuscular leads in eight arm muscles (**Figure 1A, Extended Data Figure 1**). Before SCI, we observed clear bursts of EMG activity from all hand and arm muscles during the three movement phases: reach, grasp, and pull in all monkeys. Finally, to document the involvement of cortical neurons during movement enabled by EES and to extract signals that could also be used to link stimulation bursts to movement phase onset, we implanted multi-microelectrode arrays (Blackrock Microsystems, Salt Lake City, USA) in the arm/hand region of the right sensorimotor (M1, S1) and ventral premotor (PMv) cortex. We validated these recordings by verifying that neural activity was consistently modulated with kinematics pre-injury and with the three movement phases as largely expected^54^ (**Figure 1, Extended Data Figure 1**). In summary, we analyzed natural arm movements in monkeys and concluded that in order for stimulation protocols to be effective, it was important to support reach, grasp and pull independently with specific parameters for each animal.

**Figure 1.**
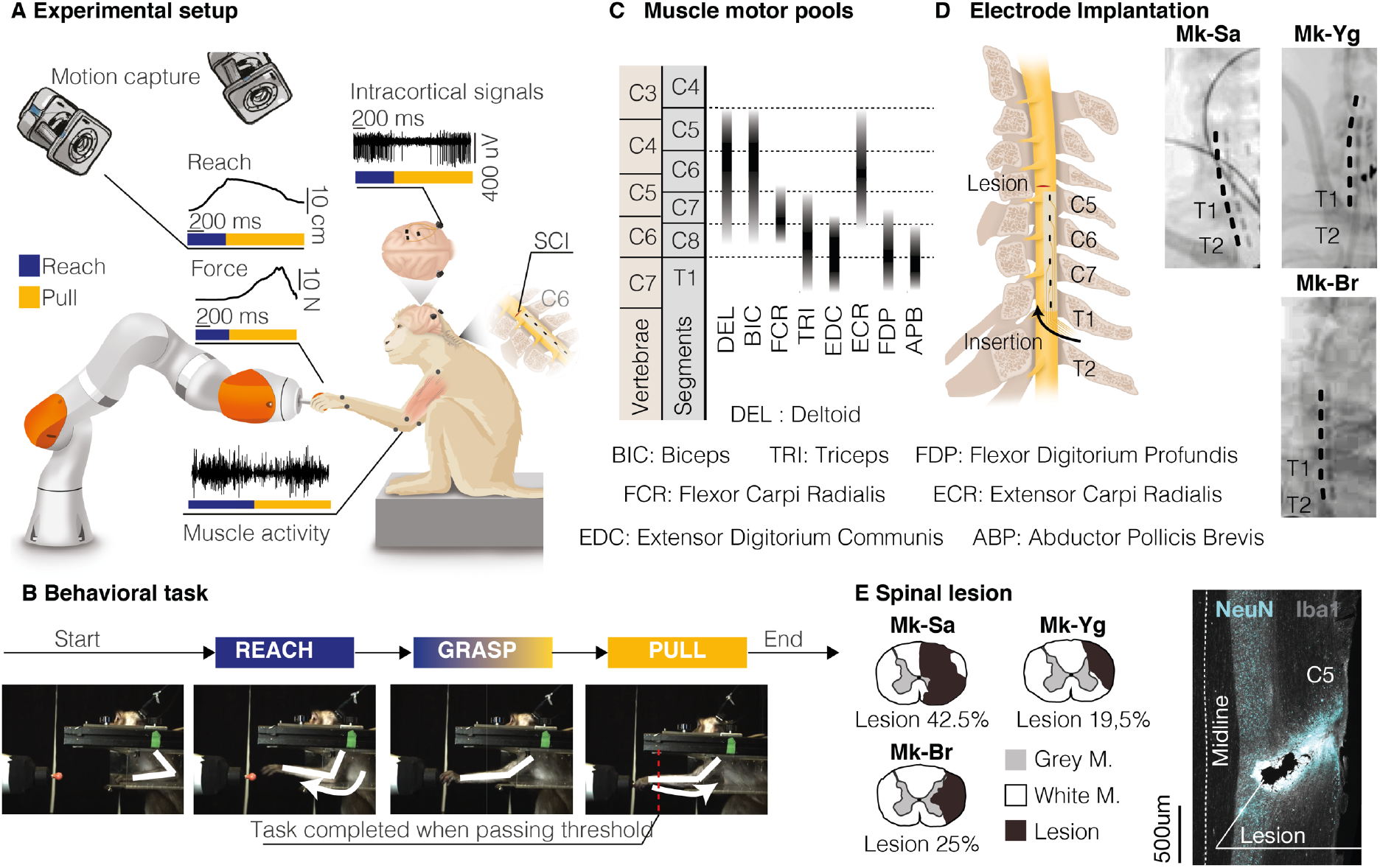
Experimental framework. **(A)** Schematic of the behavioral experimental platform. While the animals were performing a robotic reach, grasp and pull task, we measured 3D forces applied to the robot joints, full-limb kinematics, electromyographic (EMG) activity from eight muscles of the arm and hand, and intra-cortical signals from sensorimotor areas. **(B)** Schematic illustration of the task. Monkeys were trained to reach for, grasp, and pull a target object placed at the end effector of a robotic arm. We considered a movement complete when a target spatial threshold was crossed during pull. **(C)** Motoneurons pool distribution of arm and hand muscles in the cervical spinal cord in relation to vertebrae and spinal segments (adapted from Jenny and Inukai, 1983). Deltoid (DEL), Biceps Brachii (BIC), Flexor Carpi Radialis (FCR), Triceps Brachii (TRI), Extensor Digitorium Communis (EDC), Extensor Carpi Radialis (ECR), Flexor Digitorium Profundis (FDP), Abductor Pollicis Brevis (ABP). **(D)** Schematic representation of spinal implant positioning and X-ray scans of the epidural implant in the three monkeys (Mk-Sa, Mk-Br and Mk-Yg). **(E)** Anatomical reconstruction of the cervical spinal cord lesion (black area) for the 3 monkeys, shown on a transversal section (the percentage indicates the portion of the total spinal cord area that was injured on this transversal plane). On the right, representative image of longitudinal section of the spinal cord of Mk-Br around the lesion site stained with NeuN (neuronal cell bodies) and Iba1 (microglia). Copyright Jemère Ruby.

### Personalized spinal interface

To design an optimal interface, we studied the anatomy of the monkey cervical spinal cord. We extrapolated available anatomical information from literature and found that, similar to humans, motoneurons innervating arm muscles in the monkeys are segmentally organized^55^ (**Figure 1C**). We previously showed that stimulation of a single cervical dorsal root will recruit motoneurons that receive direct afferent inputs from that root^53^. Exploiting this property allows to obtain a segmental recruitment order of motoneurons that can be targeted to promote specific movement phases^49,51,56^. Therefore, we designed a spinal interface that could target each root independently. We achieved this by placing contacts on the lateral aspect of the cord to target the entry zone of each individual root^53^. Since each monkey displayed a unique anatomy, we tailored the design of our interface to each specific subject. For this, we measured white matter diameter and vertebral canal features from computed tomography (CT) and magnetic resonance imaging (MRI). We then spaced the electrodes rostro-caudally and medio-laterally to match the transversal and longitudinal dimensions of the cord of each animal (**Extended Data Figure 2A, 2B**). This allowed us to simplify the neural interface architecture by minimizing the number of contacts while maintaining high muscle recruitment specificity^57^. We then designed a surgical strategy to position the epidural interface between the C6 and T1 dorsal roots (**Figure 1D**). We performed laminectomies between the T1 and T2 vertebrae and the C5 and C6 vertebrae, then pulled the neural interface through the intermediate epidural space with the help of a custom soft inserter^57^.

We verified that the position of the array remained stable for the entire duration of the study (up to 3 weeks) through repeated X-ray imaging (**Figure 1D, Extended Data Figure 2C**). During the same surgery, we performed a unilateral spinal cord injury at the C5/C6 segments (**Figure 1E**) aiming at transecting the cortico-spinal tract that is located on the lateral aspect of the white matter in monkeys. This type of lesion is amply described in literature and induces unilateral arm and hand paralysis^58,59^ while preserving important bodily functions such as bladder control. Postmortem immunohistochemistry analysis of the spinal cords showed that the spinal interface did not damage the cervical cord in any of the three monkeys but did reveal that Mk-Br received an unplanned compression injury at the insertion site (T3 spinal segment). Given the caudal position of this contusion it is likely for it to have occurred during implantation (**Extended Data Figure 2D**). Since the T3 segment is below the innervation of the arm motoneurons, this lesion did not affect the phenotype of arm and hand motor deficits which did not differ from the other monkeys (see Methods).

In summary, we designed a spinal interface to selectively recruit the cervical dorsal roots. We tailored the interface to the specific anatomy of each monkey and designed a surgical strategy to perform a consistent and stable implantation.

### Cervical EES produces functional joint movements and grasp in anaesthetized moneys

We next assessed the selectivity of the epidural interface. In propofol anaesthetized monkeys, we delivered asymmetric, charge-balanced biphasic pulses of EES at low repetition rate (1Hz) at various current amplitudes from each contact. Minimum and maximum amplitude values were selected as the first subthreshold and first saturation current value respectively. As predicted^53^, different stimulation contacts generated muscle recruitment patterns that mirrored the segmental organization of cervical motoneurons (**Figure 2A, Extended Data Figure 3**). Specifically, contacts located at C8/T1 level (caudal) elicited spinal reflexes mostly in the hand and forearm muscles, contacts located at C7 level elicited triceps and contacts located at C5/C6 recruited biceps and deltoids (rostral). Those results were consistent in all animals (**Figure 2B, Extended Data Figure 3**). To ensure that this segmental selectivity translated into separate functional arm and hand movements, we delivered supra-threshold stimulation at various frequencies (20-120 Hz) from each contact in two animals (Mk-Br and Mk-Yg). Indeed, since recruitment of motoneuron is pre-synaptic, EES may not be able to produce sustained muscle activation because of frequency dependent suppression^60^. This effect is an observed substantial suppression of muscle evoked potentials during repetitive stimulation of the afferents. Instead, we observed large and sustained arm movements during EES bursts. Muscle selectivity was preserved during long stimulation trains (**Figure 3C, F**) and different contacts elicited distinct functional joint movements (**Figure 3A, B, D, E, Video 1**) such as shoulder abduction, elbow extension and whole hand grasp. When looking at the energy of the EMGs, we found a monotonic relationship between muscle activation and stimulation frequency in most of the upper arm muscles (**Figure 3C, F**). However, not all muscles showed such clear frequency dependent responses (**Extended Data Figure 4A**). Moreover, peak-to-peak responses (**Extended Data Figure 4B**) were generally decreased during a burst at high frequency but were not abolished and tended to vary during the burst and while the movement was produced. We used these observations to optimize stimulation parameters to be used in a behavioral reach and grasp task (see Methods and **Extended Data Figure 5**). In summary, we found that single contacts of our spinal interface elicited segmental recruitment of arm flexors, extensors and hand flexors. Bursts of stimulation from these contacts produced sustained joint movements that were graded by stimulation frequency (**Extended Data Figure 6**).

**Figure 2.**
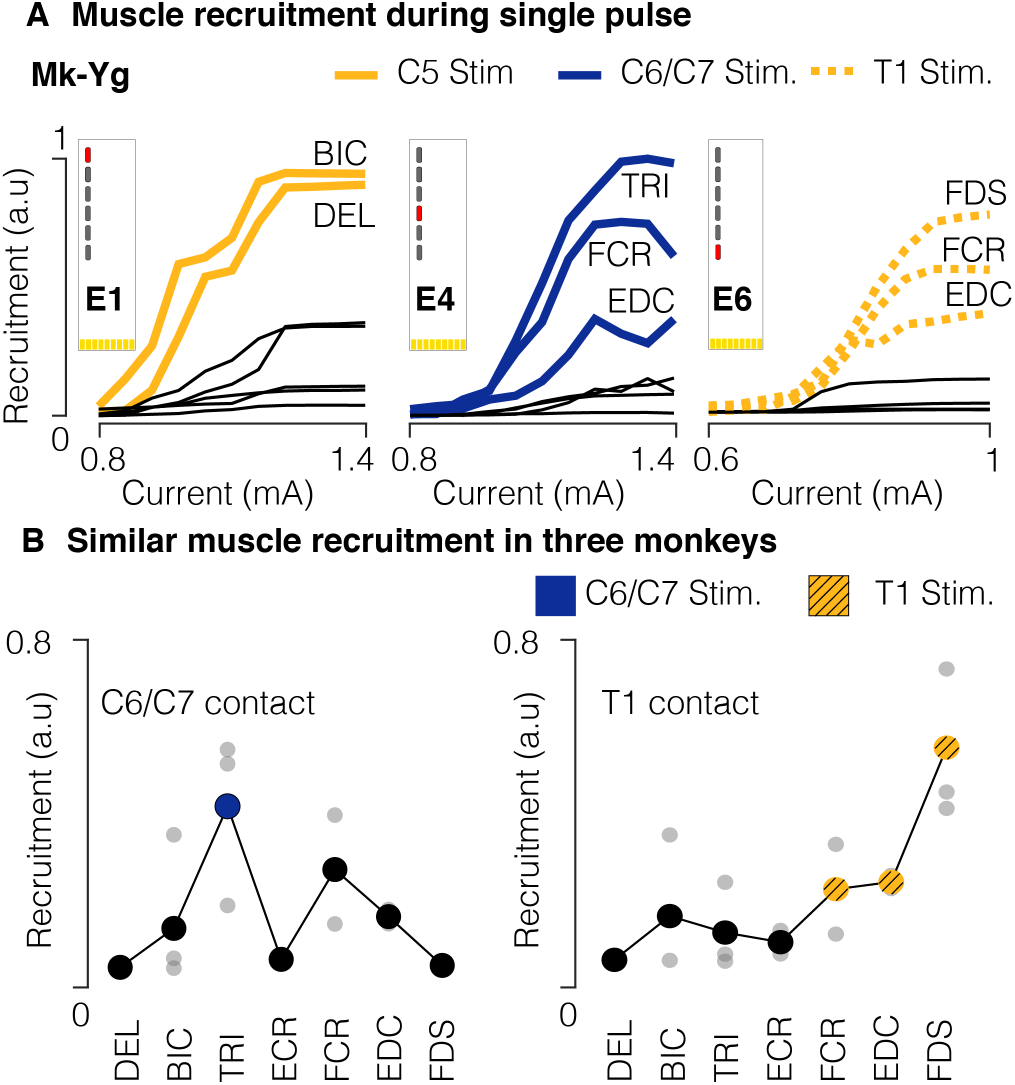
Muscle recruitment of spinal stimulation. **(A)** Examples of muscle recruitment obtained by stimulating (1 Hz) at C5, C6/C7, and T1 spinal segments (Mk-Yg). **(B)** Average muscle activations elicited from C6/C7 and T1 contacts in n=3 monkeys (grey bullets: for each animal, average recruitment across all stimulation currents. Big bullets: mean of average recruitments across animals).

**Figure 3.**
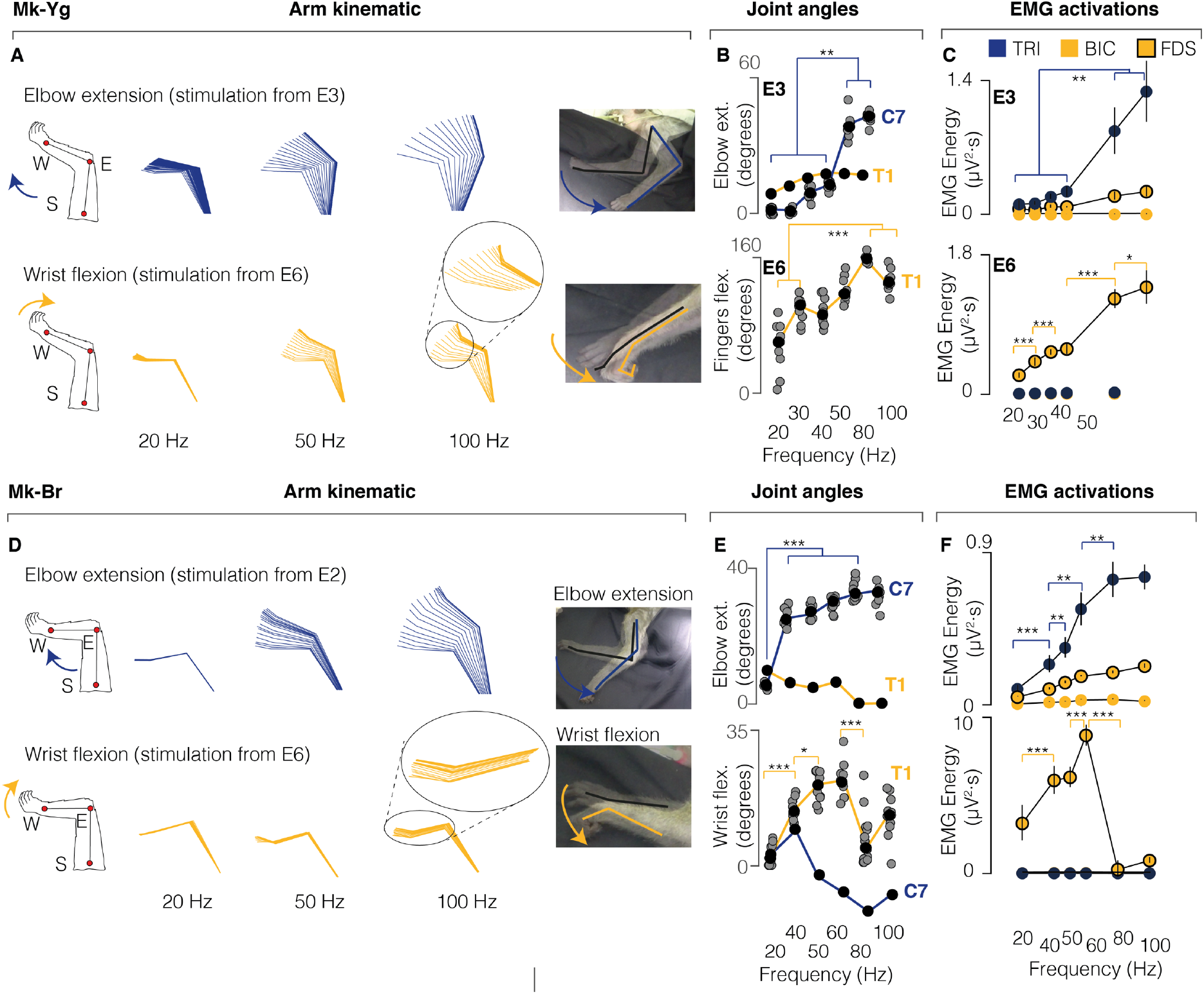
EES produces functional joint movements in anesthetized animals. **(A)** Stick diagram schematic of elbow extension and wrist flexion movements elicited by pulse-trains of stimulation in anesthetized conditions in Mk-Yg. **(B)** Modulation of maximal joint angles achieved by pulse-trains of stimulation at different frequencies, in anesthetized conditions in Mk-Yg. Stimulation was delivered at C7 (blue) and T1 (yellow). Statistics performed with Wilcoxon Ranksum test and Bonferroni correction. Asterisks: *p<0.05, **p<0.01, ***p<0.001 **(C)** Triceps (blue), biceps (yellow), and flexor digitorium superficialis (yellow with black border) activity elicited by pulse-trains of stimulation at different frequencies, in anesthetized conditions in Mk-Yg. Bullets represent mean values and bars are standard deviation. Statistics performed with Wilcoxon Ranksum test and Bonferroni correction. Asterisks: *p<0.05, **p<0.01, ***p<0.001 **(D)** Stick diagram schematic of elbow extension and wrist flexion movements elicited by pulse-trains of stimulation in anesthetized conditions in Mk-Br. Statistics performed with Wilcoxon Ranksum test and Bonferroni correction. Asterisks: *p<0.05, **p<0.01, ***p<0.001 **(E)** Modulation of maximal joint angles achieved by pulse-trains of stimulation at different frequencies, in anesthetized conditions in Mk-Br. Stimulation was delivered at C7 (blue) and T1 (yellow) **(F)** Triceps (blue), biceps (yellow), and flexor digitorium superficialis (yellow with black border) activity elicited by pulse-trains of stimulation at different frequencies, in anesthetized conditions in Mk-Br. Bullets represent mean values and bars are standard deviation. Statistics performed with Wilcoxon Ranksum test and Bonferroni correction. Asterisks: *p<0.05, **p<0.01, ***p<0.001

### Cervical EES substantially improves arm and hand motor function after spinal cord injury

We next tested whether our stimulation protocol could improve functional outcomes of upper limb movements after SCI. Specifically, we tested the efficacy of EES to improve muscle activation, pulling forces, functional task performance, and kinematic quality of three-dimensional movements after SCI when stimulation was on against stimulation off as a control. In all monkeys, the lesion led to substantial motor deficits of the left arm and hand.

While each monkey retained the ability to activate proximal shoulder and biceps muscles, elbow extension and hand functions were severely compromised. Severity of the impairment and extent of spontaneous recovery (**Extended Data Figure 7**) varied across monkeys because of the variability in lesion size (**Figure 1E**). Generally, animals showed severe paralysis immediately after lesion, and then gradually regained some movement capabilities (**Extended Data Figure 7**). Due to the initial impairment, immediately after the lesion, monkeys were not able to perform the behavioral task. Consequently, during the first week, we simplified the task by presenting an object close to the monkeys and triggering stimulation bursts manually to encourage the animal to perform the task. After the first week, all monkeys spontaneously attempted to perform the task, making it possible to link the delivery of movement-specific stimulation bursts to real-time detection of movement onset using intra-cortical signals. Whenever the monkeys strived for a reach, grasp or pull movement, we delivered bursts of stimulation promoting reach or grasp/pull respectively (movement specific EES). Outcomes were computed for each animal independently and compared between EES on and EES off. In terms of functional task performances, without stimulation, the monkeys were rarely capable of completing any part of the task (defined as reach, grasp and pull). Instead, with the support of EES, both the percentage of success and the rate of success improved with rates that depended on the level of function of the animals over time. For example, reach was recovered immediately with larger improvements at the beginning, when deficits were larger in all three animals. Instead, improvements in grasps emerged only later when the animals spontaneously recovered some movement capacity (**Figure 4, Video 2,3,4**). More specifically Mk-Br improved grasp and pull only after 2 weeks with stimulation while Mk-Yg was never able to grasp and pull except during stimulation which we could test only until day 7 when the grasp contact E6 failed (see Methods).Instead, when we used our interface to deliver continuous EES that was not related to movement onsets, only non-significant and modest improvements were observed in Mk-Br while Mk-Yg did not show ability to grasp and pull during continuous EES (**Extended Data Figure 8A**). Moreover, we analyzed trials in which stimulation bursts were not triggered at movement onset, for example when pull stimulation was erroneously triggered during reach. In these trials the reach movement was abruptly interrupted, and the animal did not complete the task (**Extended Data Figure 8B, Video 5**).

**Figure 4.**
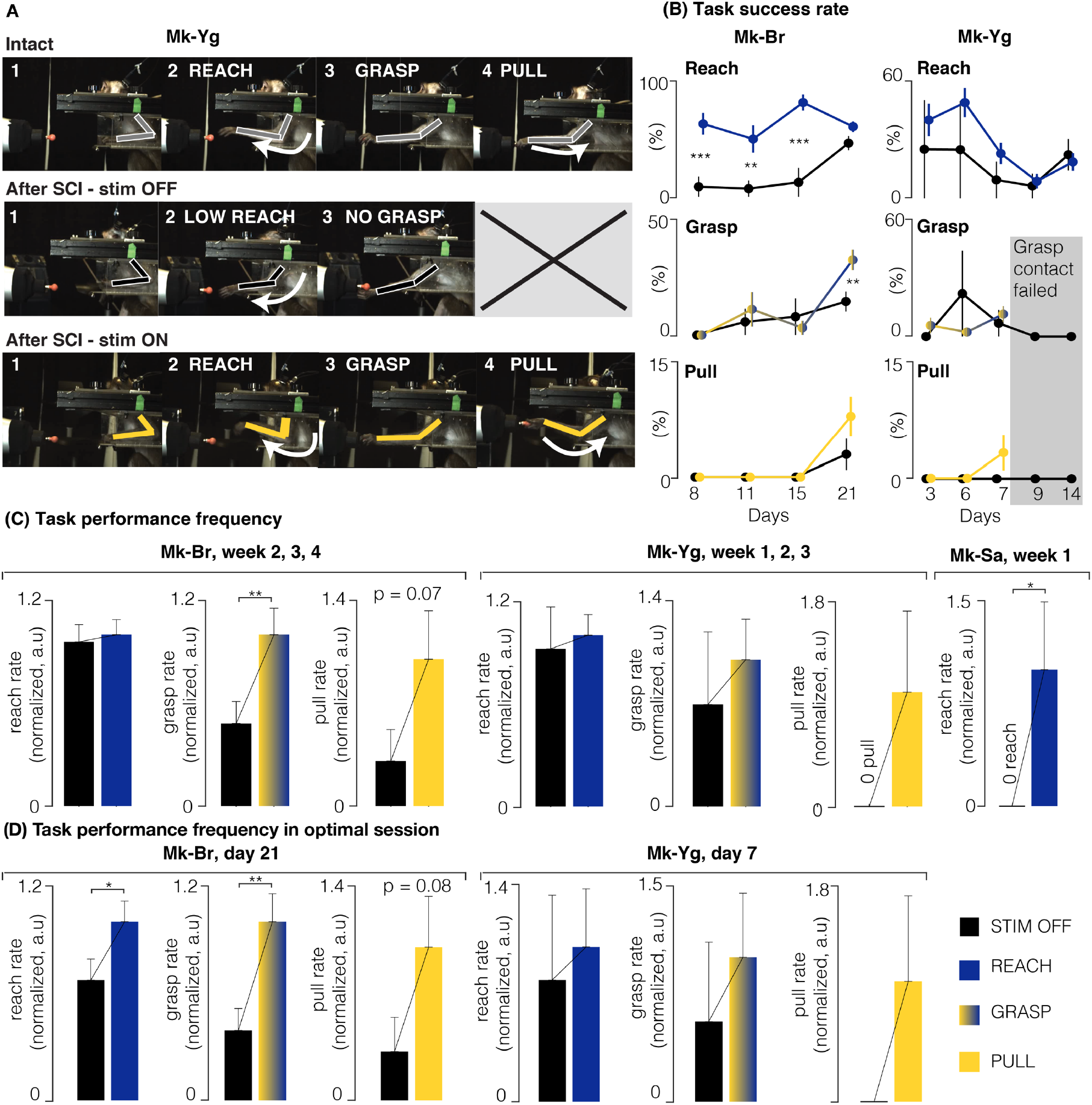
EES improves task performance. **(A)** Snapshots of Mk-Yg performing the task before SCI, after SCI without EES, and after SCI with EES. A full successful trial is composed of a reach, a grasp, and a pull. After SCI, Mk-Yg could only perform reaching movements without EES, while when EES was delivered the full task could be performed. **(B)** Task performance rate over all available sessions, computed as the percentage of successful movements across all attempted movements. Performance rate are shown for reach (blue), grasp (yellow to blue gradient) and pull (yellow movements). Data are shown as mean (bullets) and standard deviation (bars). Statistics was performed with Bootstrap. Asterisks: *p<0.05, **p<0.01, ***p<0.001. **(C)** Bar plots report the rate of successful movements after SCI without and with stimulation, for all the days in which animals performed the task. Rates were computes as number of successful trials per units of time. Data are presented as mean ± STD and normalized on the mean value in stimulation condition. Statistics was performed with Bootstrap, Asterisks: *p<0.05, **p<0.01, ***p<0.001. (P-values. Mk-Br: reach, n.s; grasp, p =0.01; pull, p =0.08. Mk-Yg: reach, grasp and pull n.s) **(D)** Bar plots report the rate of successful movements after SCI without and with stimulation, for the best session of Mk-Br and Mk-Yg. Data are presented as mean ± STD and normalized on the mean value in stimulation condition. Statistics was performed with Bootstrap, Asterisks: *p<0.05, **p<0.01, ***p<0.001. (P-values. Mk-Br: reach, p = 0.04; grasp, p = 0.002; pull, p =0.08. Mk-Yg: reach, grasp and pull n.s)

During phase dependent stimulation, EES enhanced muscles activity and forces (**Figure 5A,B**) compared to no stimulation. In terms of movement quality, EES bursts triggered at movement onset significantly improved the overall quality of arm movements (**Figure 5B**). Indeed, principal component analysis (PCA) of three-dimensional kinematic parameters (i.e., timing, force, arm trajectories, joint angles) revealed that during EES, movement kinematics were significantly closer to pre-lesion kinematics than the few successful movements performed without stimulation (distance from pre-lesion performances in the multi-parametric kinematic space, **Figure 5B**). Notably, animals sustained the weight of the arm and lifted their elbow more, performed wider movements, and generated stronger forces (**Figure 5B**), getting closer to normal kinematic trajectory patterns without any long-term training.

**Figure 5.**
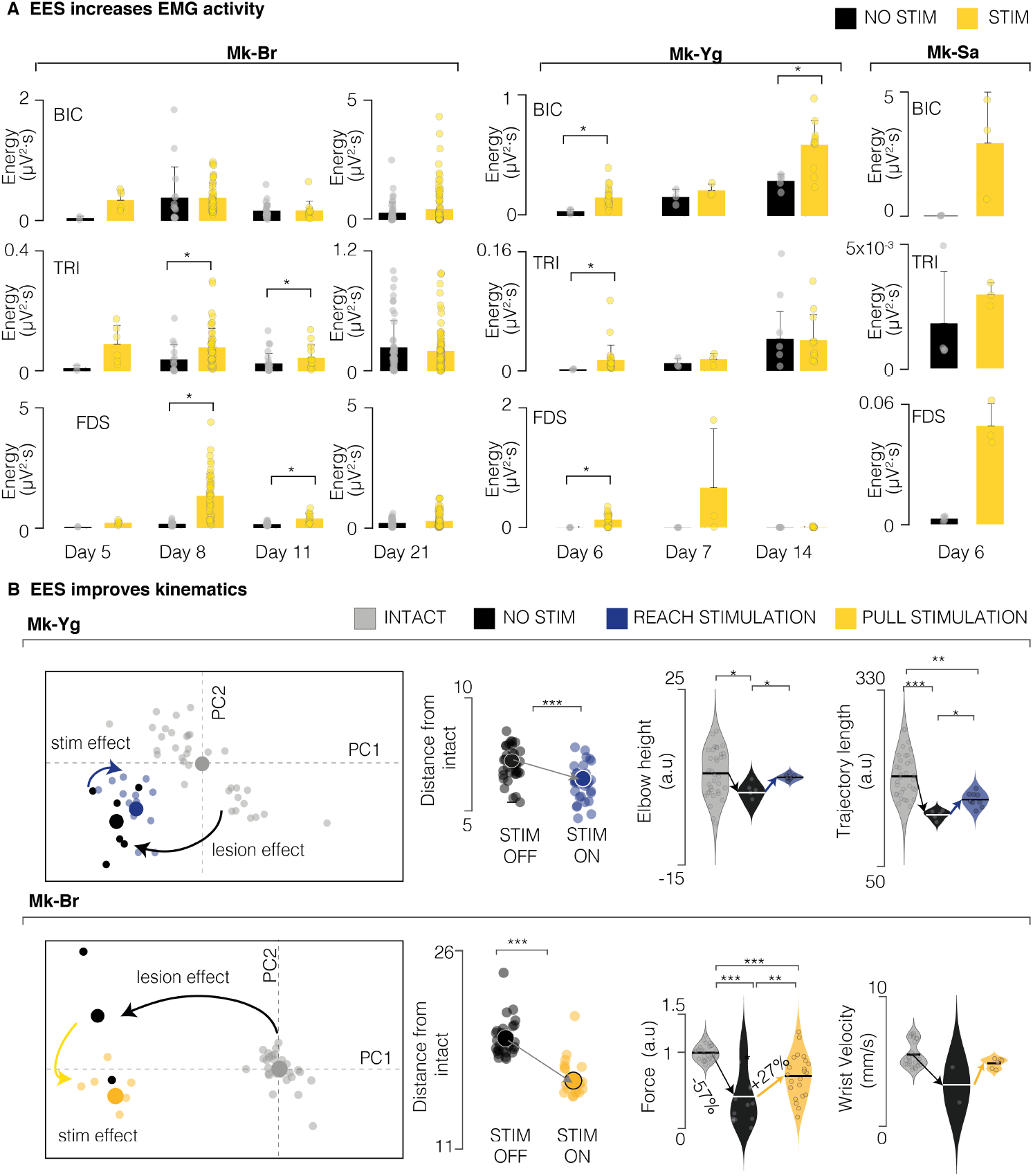
EES improves muscle strength and movement quality. **(A)** Bar plots of signal energy of biceps, triceps and FDS EMG profiles during movement with no stimulation (black) and stimulation (yellow). Data are shown for different sessions (one for each week) in Mk-Br and Mk-Yg. Mk-Sa performed only one session. All individual data points are represented by bullets. Statistical analysis with Wilcoxon Ranksum test. **(B)** PC analysis of kinematic features for Mk-Yg (top) and Mk-Br (bottom). From left to right: (1) first and second PC space. Each bullet represents one trial. Trials performed after injury (black) are consistently separated from the trials performed in intact conditions, highlighting a change in the quality of resulting kinematics. Trials performed with the support of stimulation (blue for reach and yellow for pull) are located closer to the intact trials in the PC space, denoting an improvement in kinematic features. (2) euclidean distance in the feature space of trials without stimulation (black) and with stimulation (blue for Mk-Yg, yellow for Mk-Br) from the centroid of the trials in intact condition; (3) example violin plots of movement quality features in the three conditions: intact, after SCI, and after SCI with stimulation. Statistics with Wilcoxon Ranksum test. Asterisks: *p<0.05, **p<0.01, ***p<0.001.

In summary, we showed that EES bursts triggered at movement phase onsets, improved muscle strength, task performance and quality of arm movements. This allowed monkeys to perform reach, grasp and pull movements that were otherwise not able to perform without EES.

### Sensory inputs can decrease EES-induced motor output

We then investigated the role of spinal circuits and sensory inputs in the production of the movements that we observed. Indeed, since activation of motoneurons was pre-synaptic, spinal reflexes and sensory inputs can influence EES evoked spinal reflexes in the legs^22,61^. In order to exclude influences of residual supraspinal voluntary inputs, we conducted experiments under propofol anesthesia (**Figure 6A**) with Mk-Br. We then delivered bursts of EES from the contact eliciting elbow flexion at varying stimulation frequencies in two distinct conditions (**Figure 6B**): in isometric and unconstrained conditions. In the isometric condition, we constrained the wrist, elbow and shoulder of the animal and measured force production at the wrist joint. Under unconstrained conditions we left the arm free to move under the effect of stimulation. This setup only differs from the sensory feedback generated at the load when pull forces are produced by EES. We found that EES induced EMG activity during unconstrained movement that was significantly different from the EMG activity induced during isometric movements (**Figure 6B**). In particular, overall EMGs and peak-to-peak amplitudes of elicited spinal reflexes were significantly lower when the arm was attached to a load (isometric) compared to when it was free to move. Albeit present at all frequencies, this difference was particularly important within the 40 to 60Hz range, thus overlapping with the functional frequency ranged that we selected for our study.

**Figure 6.**
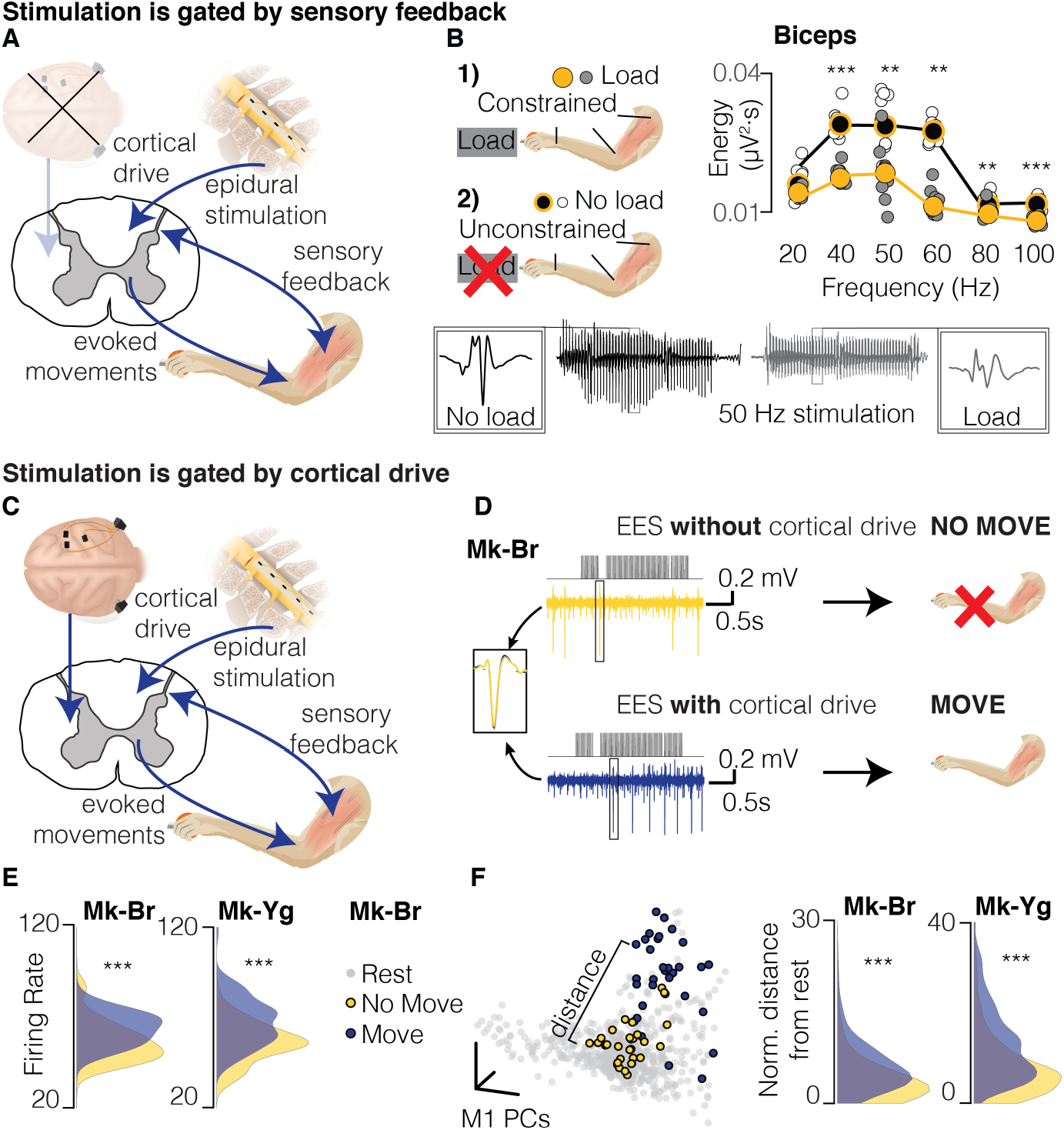
EES must be synchronized with motor intention. **(A)** Schematic of the interactions between EES and residual neural structures during anesthetized stimulation. During anesthesia, cortical control has no interaction, therefore EES interacts solely with sensory feedback spinal circuits. **(B)** Quantification of EMG activity during EES in two conditions: unconstrained arm (no load, black); arm constrained by load applied at the hand (load, gray). White and grey bullets: individual data points for no load and load conditions. Black and yellow bullets: mean values for no load and load conditions. Black and yellow lines: interpolation of mean values for no load and load conditions. On the bottom, example of EMG traces obtained during stimulation in the no-load (black) and load (gray) conditions. Stimulation artifacts have been removed. Data from Mk-Br **(C)** Schematic of interactions between EES and residual neural structures during the performance of the behavioral task. EES interacts with descending cortical drive sent through residual pathways after SCI, as well as with sensory spinal circuits. **(D)** Schematic illustrating the kinematic outcome of the interaction between EES and residual cortical inputs. The same EES pulse train (top) applied to Mk-Br can result in different motor outputs: no movement output when the cortex is silent (yellow, top), movement is produced when the cortex is active (blue, bottom). **(E)** Distribution of average firing rates across all M1 channels during stimulation trains that evoked no movement (yellow) and movement (blue). **(F)** Left: State space view of M1 activity for all time points during rest (gray), successful stimulation (blue) and unsuccessful stimulation (yellow). The brain states during unsuccessful stimulation (yellow) overlapped with the rest states, while the successful stimulation (blue) did not. Right: we computed a relative Mahalanobis distance between the two stimulation conditions and the cluster of neural states at rest. For both monkeys, neural states during stimulation periods with no movement were close to rest.

These results show that force loads at the hand decreased EMG activity induced by EES as compared to no load applied at the hand. Under anesthesia, only changes on spinal circuit excitability induced by sensory inputs can explain the observed changes on EES evoked muscle activity.

### Some residual cortical input is necessary for cervical EES to be effective

The influence of spinal sensory inputs showed that EES output may be decreased because of spinal sensory inputs when loads are applied at the hand. This would decrease the efficacy of EES which is supposed to enhance force production. Therefore, to explain the results we obtained in behaving monkeys (**Figure 6**) we investigated the contribution of residual cortical inputs in the production of forces and movements during EES. Specifically, since cortical inputs actively modulate spinal circuits, they should be able to both enhance and suppress EES output by modulating spinal circuit excitability^30^. Since we showed that monkeys could use EES to amplify their movement and forces (**Figure 6D**) we focused on demonstrating that cortical inputs could also suppress unwanted EES-generated movements. We hypothesized that if monkeys did not want to move, EES would not produce the large joint movements that we observed when the monkeys were anesthetized. Therefore, we identified trials in which our decoder detected a false-positive reach movement (**Figure 6C**). In this situation our system would deliver a burst of stimulation even if the animal was not attempting to execute the task. We then compared intracortical activity from the primary motor cortex (M1) of Mk-Br and Mk-Yg during these false-positive trials to the signals recorded during correctly detected trials. We identified trials where EES was present and the monkey moved, and trials when EES was present but the monkey did not move (**Figure 6D**). We verified that the same neural units were present in both conditions and found that the overall firing rates of all units in motor cortex was significantly higher when EES produced movement (**Figure 6E**) than when it did not. This suggested that movement happened only if the motor cortex was active, despite EES was delivered at amplitudes that generated large joint movements when the same monkey was anesthetized. To further validate this hypothesis we applied dimensionality reduction using Principal Component Analysis to the firing rates in each electrode and reduced the M1 population activity to low-dimensional states^62^. In this low-dimensional space each point represents the global neural state of the motor cortex at a given time point (**Figure 6F**). We compared the neural states present when EES was associated movements and those when EES was not associated movement with the neural states associated to rest, e.g. when the monkeys were resting before the go signals between trial repetitions. When looking at the spatial distribution of neural states, trials in which EES was not associated to movement seemed to overlap with states of rest. We then computed the distance between each neural state to the subspace representing neural states at rest and found that the neural states associated to movements during EES were significantly further away from neural states at rest than neural states associated to EES and no movement. In summary, we found that the motor cortex activity was similar to the activity at rest whenever we delivered EES but the monkey did not move (**Figure 6F**). Instead, the monkey moved when the motor cortex was significantly active. This implies that the residual cortical inputs via direct and indirect pathway can either suppress or enable movement during EES.

## Discussion

We showed that EES of cervical spinal cord immediately enhanced muscle activation and strength, task performances and movement quality during a natural-like reach and grasp task in monkeys with unilateral cervical SCI compared to no stimulation controls in three monkeys. Importantly, our technique allowed monkeys to support the weight of their arm during reach, grasp and pull movements. These results are important in light of clinical translation of our technology. Stronger forces and better arm weight bearing can empower patients with the capacity to perform a larger spectrum of movements than they would normally be capable of doing without the need of support. This may provide for more independence in daily living as well as better outcomes of physical therapy.

### Exploiting subject-specific anatomy to simplify technology

We obtained our results with relatively simple stimulation protocols that engaged up to three monopolar contacts (one for reach, one for grasp and one for pull). The combination of simple bursts through these contacts enabled whole arm multi-joint movements. We believe that the design of our interface was key to achieve this result. The dorsal roots are a robust anatomical target that we could easily identify through standard imaging to personalize surgical planning and interface design. A similar surgical planning approach can be imagined in humans where MRIs and CT can guide surgical planning^51,63^.

Our results were enabled by the relative mapping between each dorsal root and the rostro-caudal distribution of motoneurons in the cervical spinal cord, which is similar in monkeys and humans^53,55,64^. The anatomical separation of roots in the cervical enlargement allowed us to recruit each root independently which generated distinct joint movements to a degree that was not observed in applications of EES for the lower limbs^49^. Stimulation of the C6 root elicited distinct arm flexion, C7 stimulation produced arm extension and C8/T1 stimulation produced hand grasp. However, similarly to other spinal cord stimulation studies we could not identify contacts that selectively produced finger extension^18,65,66^. This is likely caused by the overlap of extensor motor-pools in the forearm^55,64^ but possibly also because flexors may be biomechanically stronger and dominate hand kinematics in the case of co-contraction at rest. Despite these limitations in specificity, we were able to restore a whole three-dimensional arm movement by solely detecting movement onset signals to trigger pre-determined stimulation bursts through two or three contacts. Unlike FES, this is possible because EES activates cervical motoneurons via pre-synaptic inputs thus allowing modulation of elicited muscle responses that can compensate for reduced specificity^30,49^.

### Supporting arm movement phases independently

Contrary to previous pilot applications of epidural and transcutaneous spinal cord stimulation of the cervical spinal cord^35,36^, we utilized a soft epidural interface that allowed selective and independent support of each movement phase rather than providing continuous stimulation to the whole spinal cord. This approach is not possible with transcutaneous technologies^67^ or current design of human leads^53^ and would require new interfaces designed for the cervical cord. Selective spatiotemporal stimulation was shown to be more effective in animal models and humans than continuous stimulation in the sense that it was able to immediately produce coordinated locomotion compared to continuous stimulation that instead required long training periods^28,48,49,49,56^. In the case of the upper limb we believe that this approach was critical. Indeed, while continuous stimulation did provide some level of facilitation, it failed to entirely promote grasp and pull in one of the monkeys. Perhaps the intrinsically unstructured nature of arm and hand control makes a continuous stimulation approach less effective than it is in locomotion that instead has an intrinsic repetitive structure^38^. For example, stimulation parameters that promote grasp, may impair reach if they are delivered continuously throughout movement. Indeed, when a pull stimulation was triggered at mid-reach it generated the interruption of the reach movement. Perhaps a different interface design or lower stimulation amplitudes could be used to optimize continuous stimulation protocols, but it would be at the expense of power of elicited movements potentially preventing the weight bearing component necessary for three-dimensional movements. In summary, the complex articulation of arm and hand movements may exacerbate the difference in efficacy between continuous and phase-specific stimulation protocols that was already observed for EES in locomotion, possibly explaining the difference in effect size that was obtained so far for application in the upper limb.

### The role of sensory feedback and residual cortical inputs in cervical EES

We showed that sensory feedback when the hand was constrained to a force load reduced the EMG power produced by EES compared to free movements. This is likely caused by afferent inhibitory feedback coming from Ib afferents^68^. Unfortunately, lower muscle power while resisting a force load would decrease the clinical usability of this technology. We believe that this phenomenon is particularly relevant for the upper limb. Indeed, also during EES of the lumbosacral cord, the EES motor output is influenced by sensory inputs^22,61^, however sensory inputs are instrumental for locomotion and heavily contribute to the generation of the repetitive movement patterns that are required to walk^16,22,23,38,69^. Therefore, in the case of locomotion these inputs amplified and sustained EES-induced activity^16,22,23,28^. Instead arm and hand movements are produced by an unstructured sequence of primitive movements^41^ and reflexes^45^ in parallel with a sophisticated gating of sensory inputs through mechanisms such as pre-synaptic inhibition^8,70^. Therefore, residual cortical inputs become instrumental to obtain arm and hand movement with EES as shown by our analysis of intra-cortical signals during the production of movement combined with the observation that functional grasp was achieved only when the animals had recovered some level of function. Indeed, our lesions were non-complete and while most of the cortico-spinal tract was transected, multiple residual descending pathways were spared. These indirect inputs could have been used by the animals to mediate the inputs required to integrate EES and sensory inputs to produce voluntary movements. In summary, we believe that even during phase-specific EES residual cortical inputs play a critical role in enabling arm movement for cervical EES.

### Clinical significance and challenges

The most important challenge for clinical translation of EES to humans concerns the role of residual inputs. Our data show that some level of residual inputs and of function is required to enable movement, first because in awake animals EES did not initiate movements, and second because it lacks the selectivity to achieve selective finger activity. However, previous studies showed that even completely paralyzed subjects retain residual but functionally silent descending inputs^25,32,51^. Therefore, while overall efficacy may depend on injury severity, even severely injured patients may obtain benefits from cervical EES. After a period of physical training combined with EES^71^ these subjects may be able to use EES to achieve simple but functional grasp. Alternatively, more selective technologies targeting hand muscles such as FES could be combined with EES to obtain powerful yet selective movements.

The adaptation of EMG output to stimulation frequency that we observed in consequence of pre-synaptic activation of motoneurons may lead to a reduction in efficacy during long-term clinical use. Additionally, stimulation of afferent fibers may cause uncontrolled reflexes which may affect function. While we did not observe these phenomena in our data, this may be due to the relatively small size of the lesion compared to severe contusion in humans. However, data in humans with SCI suggest that stimulation protocols can be adapted to be functional even in subjects with chronic severe thoracic lesions^32,51^, therefore we expect that this will be the case also for cervical lesions. At any rate both risks can be reduced by accurate stimulation tuning and real-time adaptation of stimulation patterns^22,24,72^.

Concerning complexity of our system, in our study we detected movement onsets from intracortical activity which may be seen as a limitation for a realistic implementation of our protocol in clinical settings. However, given the simplicity of our protocol which is essentially constituted by alternation of pre-defined bursts, brain recordings may not be required in clinics. Indeed, most patients suffer from a severe but incomplete paralysis^51,73^, which spares some residual muscle activity in few muscles. While this residual activity is not sufficient to produce functional movements, it can be reliably detected and used to trigger stimulation bursts with standard clinical technologies^49,51^. In summary, we believe that by exploiting the functionality of residual spinal circuits and supra-spinal inputs, cervical EES constitutes a simple yet robust approach to the restoration of arm motor control with significant translational potential.

## Acknowledgements

The authors would like to thank Jacques Maillard and Laurent Bossy for the care provided to the animals, Dr Eric Schmidlin and Dr Simon Borgognon for their help with anaesthesia and surgery preparations, Dr Marion Badi for her help and advice during experiment preparations and experimental procedures, Dr. Andrina Zbinden for her contribution to the health survey of the monkeys, André Gaillard and Andrea Francovich for their help with the implementation of the hardware and the students of the University of Fribourg Amélie Jeanneret, Alen Jelusic, Laora Marie Jacquemet and Samia Borra for their help in processing data.

## Author Contributions

MC, BB and SC conceived the study; BB, MGP, and TM designed and implemented the hardware and software tools; SC designed the behavioral task and training strategy; GS and SL designed and manufactured the implantable interface; BB, SC, MGP and MC conducted the experiments; BB, SC, MGP and KZ performed the data analysis; SC, MD and MK trained the animals; SC, KG, NJ and QB processed the histological data; JB, GC and MC designed surgical implantation strategies and stimulation strategies. GC and JB, performed surgical implantations and lesions. EMR and MC implemented and supervised procedures on monkeys; MC, BB, SC and MGP wrote the manuscript; all authors edited the manuscript; SL, TM, JB, GC and MC secured funding for the study; MC supervised the study.

## Data and materials availability

All software and data will be available upon reasonable request to the corresponding author.

---

**Extended Data Figure 1.**
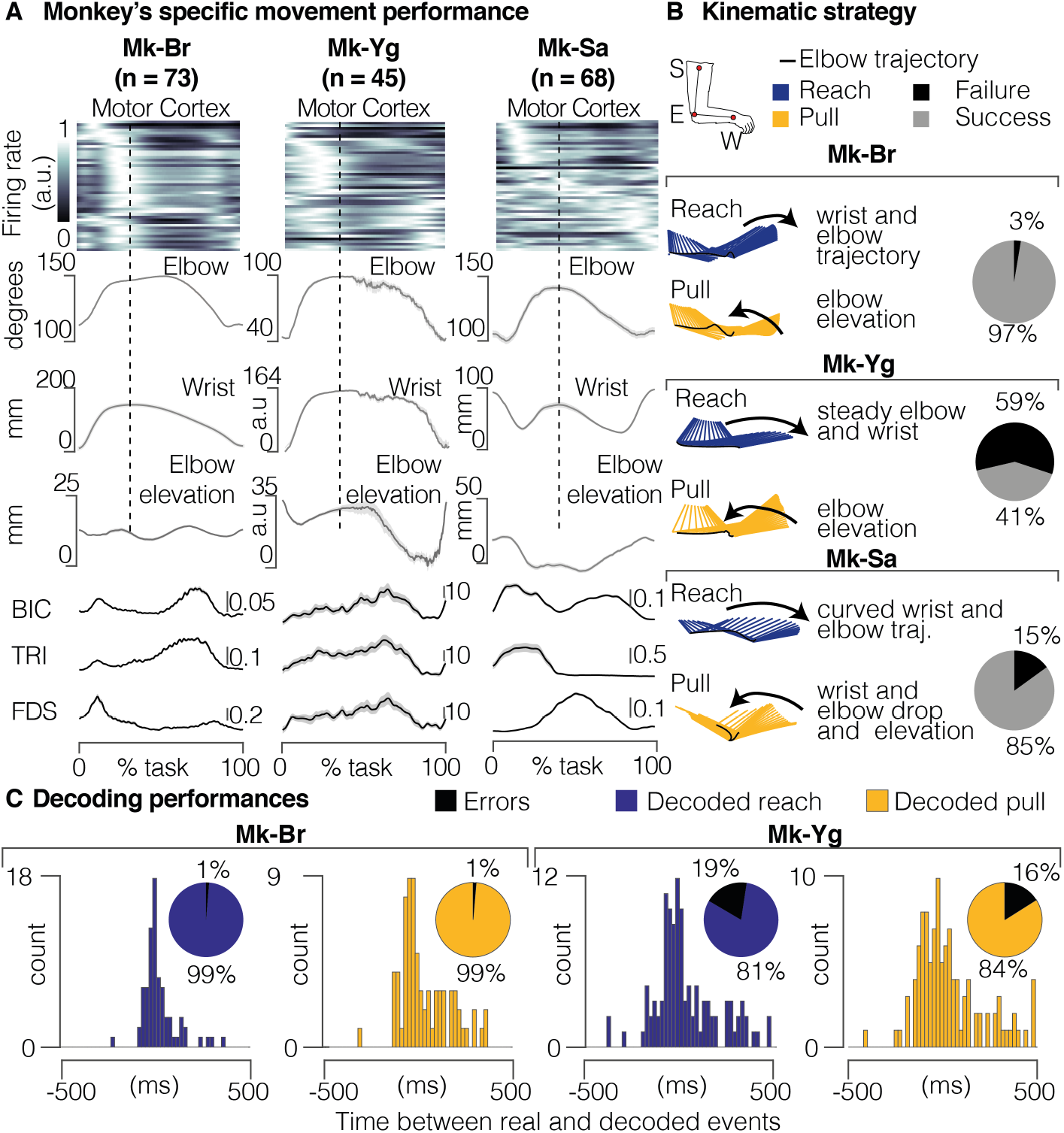
**(A)** Portfolio of signals recorded during intact movement for each animal. These signals have been recorded during the experimental session prior to the lesion. Motor cortex recordings show firing rate profiles for the 64 microelectrodes. Each row shows the firing rate of a specific electrode. Electrodes are displayed from top to bottom by order of first activation in a reference trial. Arbitrary units in motor cortex recording indicate normalized firing rate for each electrode (see methods). In kinematic and EMG plots, black lines correspond to the mean profile across all trials, shaded area shows the SEM across all trials. Kinematic scales are expressed in mm. For Mk-Yg, arbitrary units on kinematic plots represent displacement units derived by the count of video pixels. EMG scales are expressed in mV. **(B)** Kinematic strategies implemented by each monkey. Stick diagrams representations of the arm kinematic during reach (blue) and pull (yellow). The black line highlights the elbow trajectory. Pie charts represent the percentage of success and failure in task performance before lesion. **(C)** Offline decoding performance for Mk-Br and Mk-Yg before lesion. Histograms show timing accuracy of reach (blue) and pull (yellow) event decoding. The height of bars (y coordinate) illustrates the amount of events decoded with a specific timing accuracy (x coordinate). Pie charts (inset) show the percentage of correctly identified (true positive) reaches (blue) and pulls (yellow), across all decoded events. The black portion of the pie chart highlights the percentage of false positive decoded events.

**Extended Data Figure 2.**
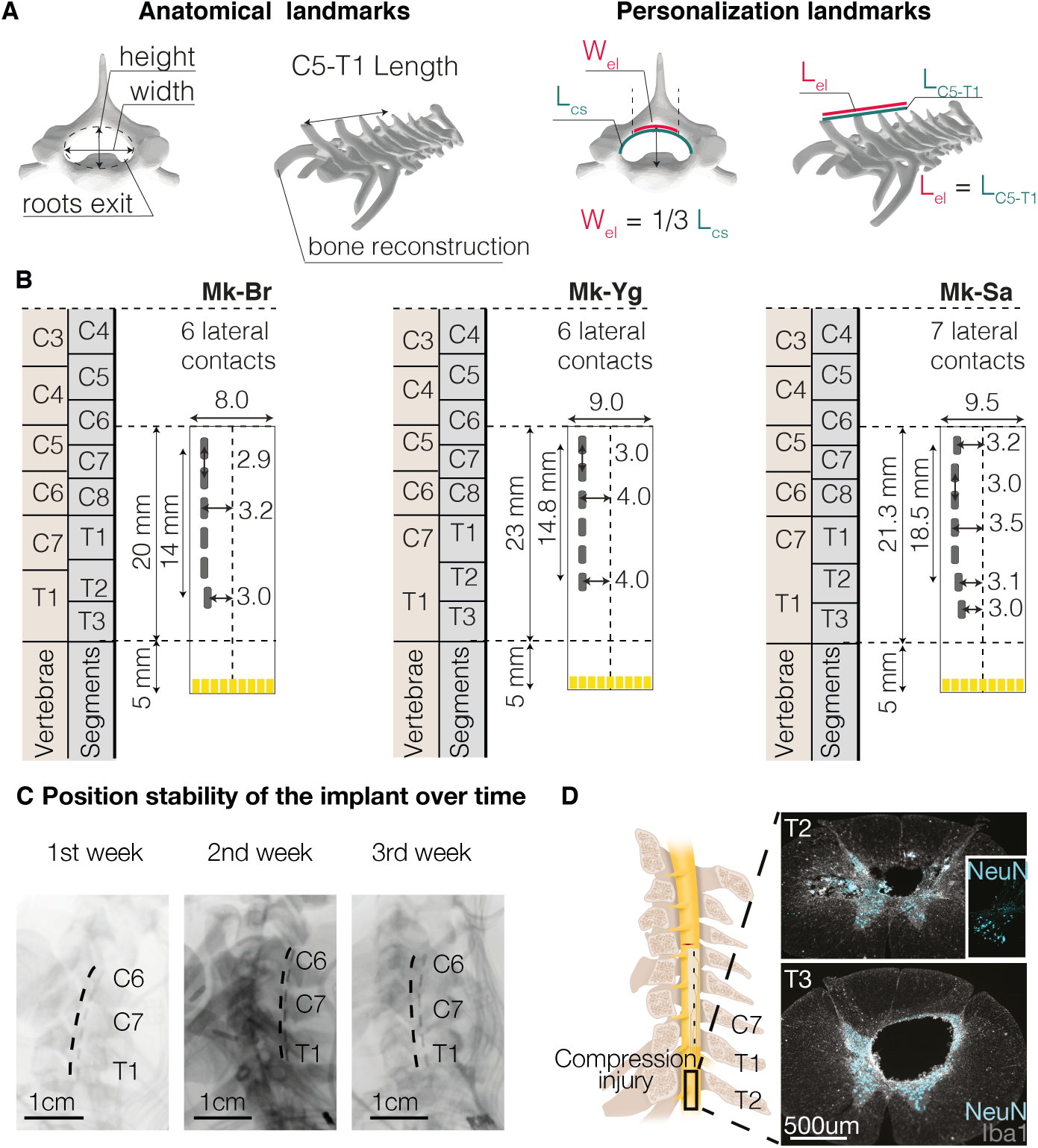
**(A)** Anatomical landmarks used to tailor the epidural interface to each monkey’s anatomy (Length of dorsal aspect of spinal canal L_CS_, length of C5-T1 spinal segment L_C5-T1_, electrode width W_el_, electrode length L_el_). Three-dimensional reconstructions of vertebras are obtained by CT-reconstruction (Osirix, Pixmeo, Switzerland). **(B)** Personalized design of the epidural implant for each animal. All measures are in millimeters. Yellow traces at the bottom of the electrode identify connectors. **(C)** Position stability of the epidural array over time, illustrated through X-rays imaging taken during 3 consecutive weeks after the implantation, images from Mk-Yg **(D)** Compression injury at the insertion level of the array (T2-T3 segment) in Mk-Br, discovered post-mortem, stained with NeuN (neuronal cell bodies) and Iba1 (microglia).

**Extended Data Figure 3.**
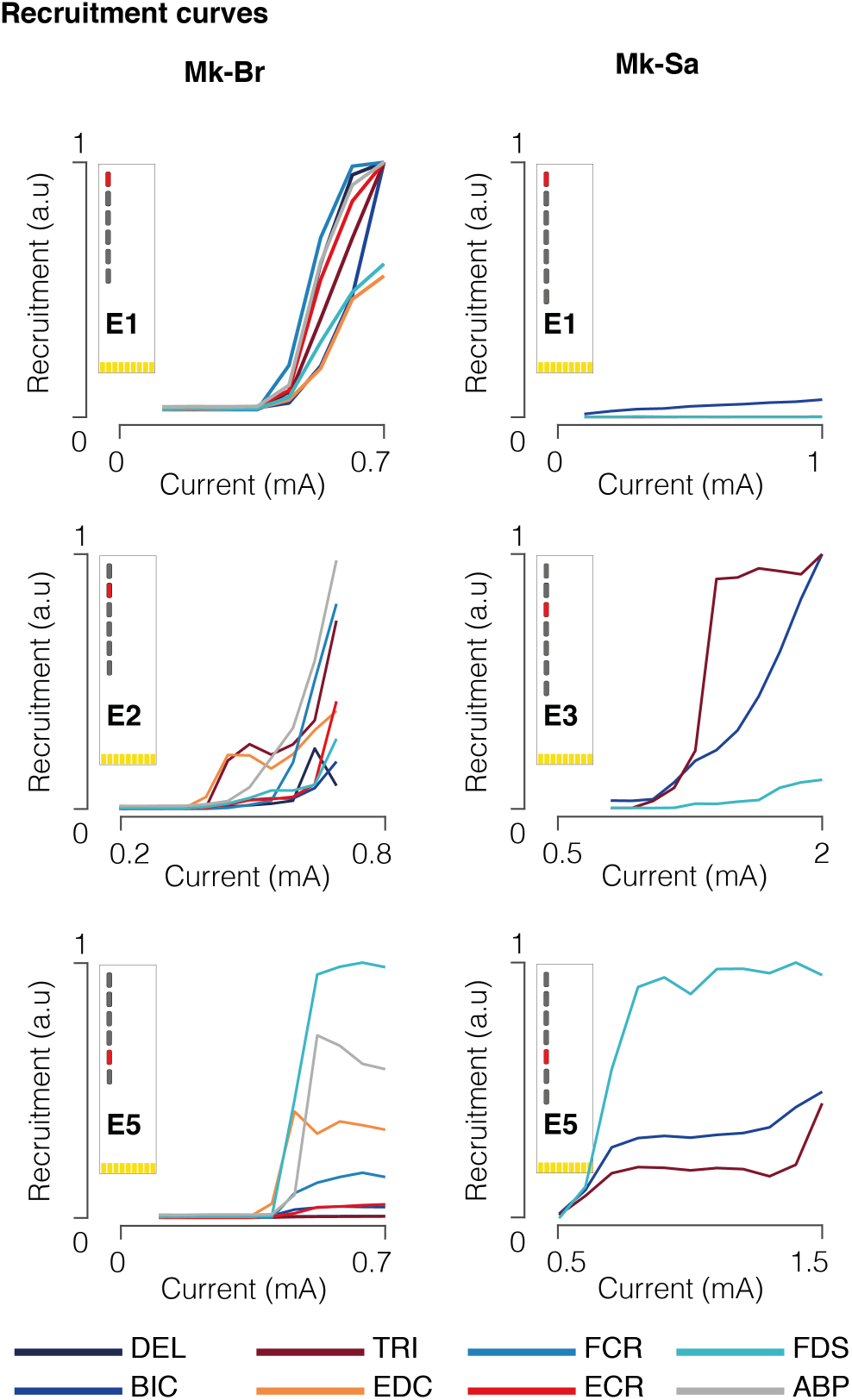
Muscle recruitment obtained by stimulating (1 Hz) at C5, C6/C7, and T1 spinal segments for Mk-Br and Mk-Sa. Mk-Sa only had three muscles implanted: biceps, triceps, and flexor digitorium superficialis.

**Extended Data Figure 4.**
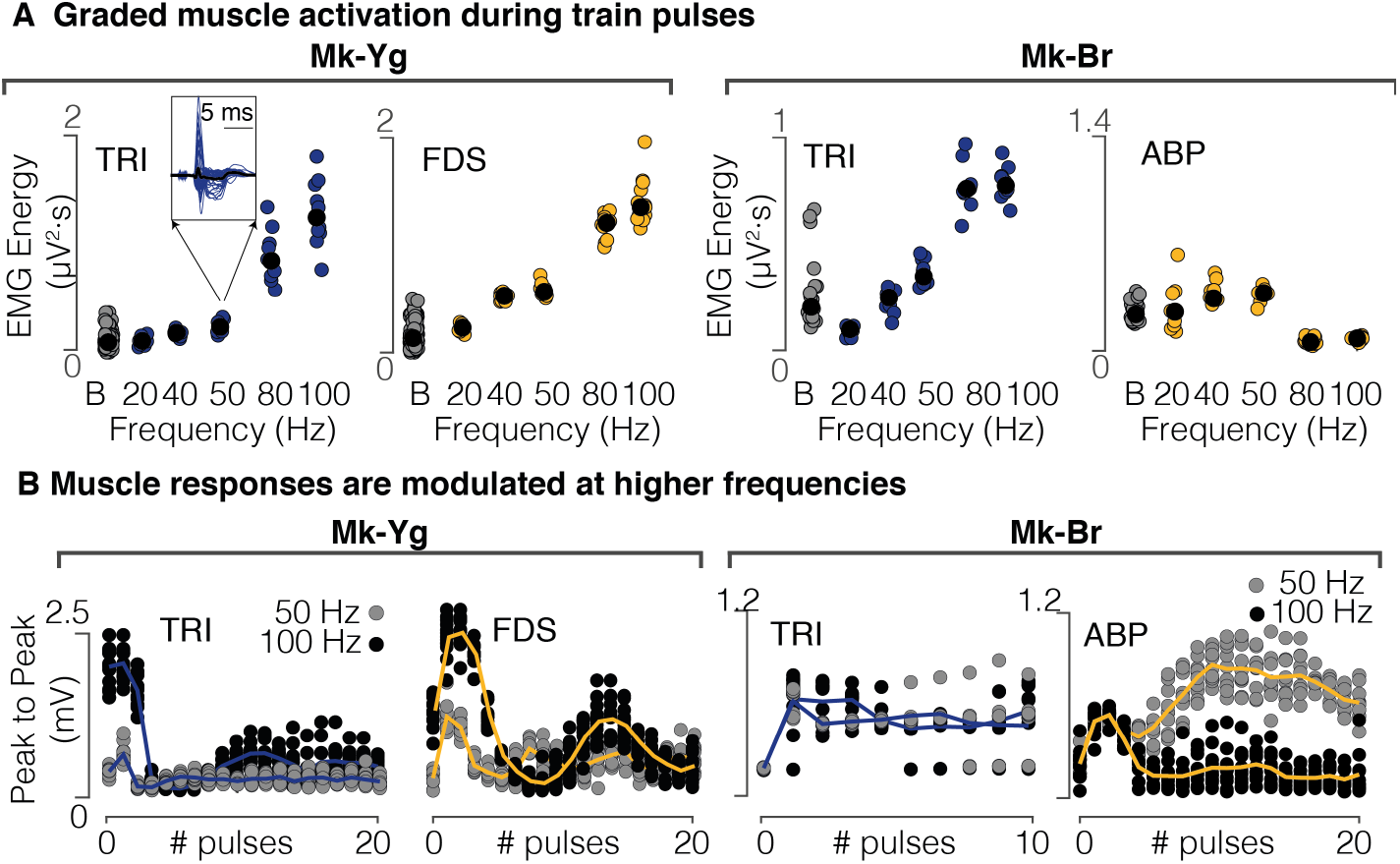
**(A)** Energy of EMG signals of triceps (Mk-Br and Mk-Yg), Flexor Digitorium Superficialis (Mk-Yg) and abductor pollicis (Mk-Br) muscles, following pulse-train stimulation at different frequencies (on the x-axis). Black bullets represent mean values. **(B)** Evolution over time of the peak-to-peak value of stimulation evoked responses during a stimulation burst. Each plot shows the evolution for a specific muscle following pulse-train stimulation at 50 and 100Hz. Triceps is shown for Mk-Br and Mk-Yg, Flexor Digitorium Superficialis for Mk-Yg and abductor pollicis for Mk-Br. Each data point is represented as a bullet and lines represent mean values over time.

**Extended Data Figure 5.**
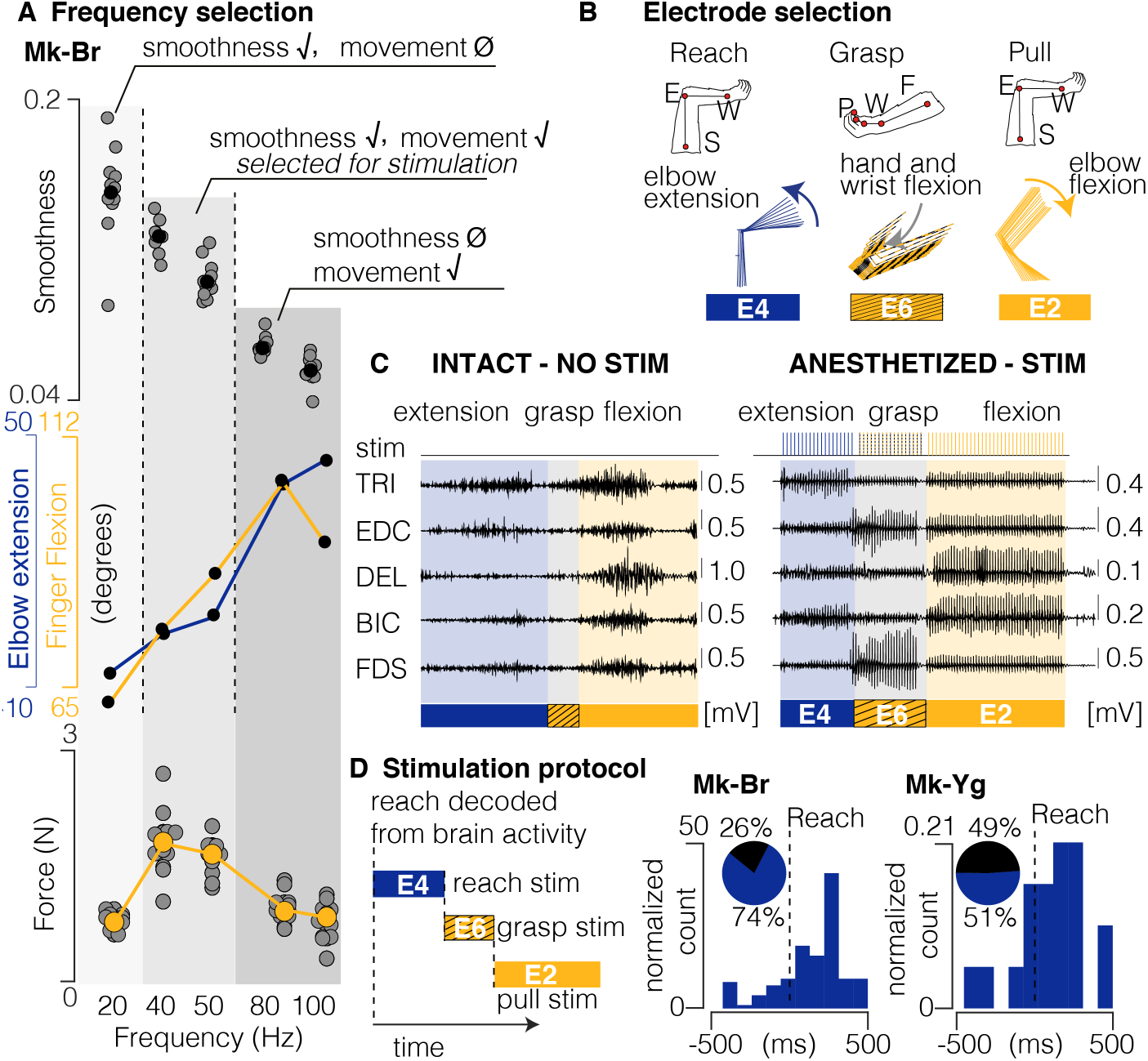
Design of stimulation protocol. **(A)** Combined representation of movement smoothness, elbow and finger flexion, and pulling force during anesthetized stimulation. Shades of gray highlight three frequency ranges that produce: (1) smooth trajectory, but little movement and low force (20Hz), (2) smooth trajectory, extended movement and medium force (40 and 50Hz), (3) abrupt and very extended movement and low force (80 and 100Hz). Kinematics and force reported here were measured in different experiments, kinematics was unconstrained, force data were acquired in isometric conditions (see Methods). The range 40-50 Hz was selected as the best optimization of sufficient movement, smoothness and force production. **(B)** Schematic representation of arm and hand kinematics during stimulation delivered from the selection of three contacts to produce elbow extension (blue), hand and wrist flexion (yellow and black), and elbow flexion (yellow). **(C)** Example of comparison between EMG activity during intact movement (left) and movement elicited by chaining stimulation from the three selected contacts (right). **(D)** Scheme illustrating how stimulation is triggered from movement-related intra-cortical signals. On the right, online performances of movement attempt decoder in two animals with SCI. Pie charts represent percentage of predicted (blue) and unpredicted (black) reach events by our decoder.

**Extended Data Figure 6.**
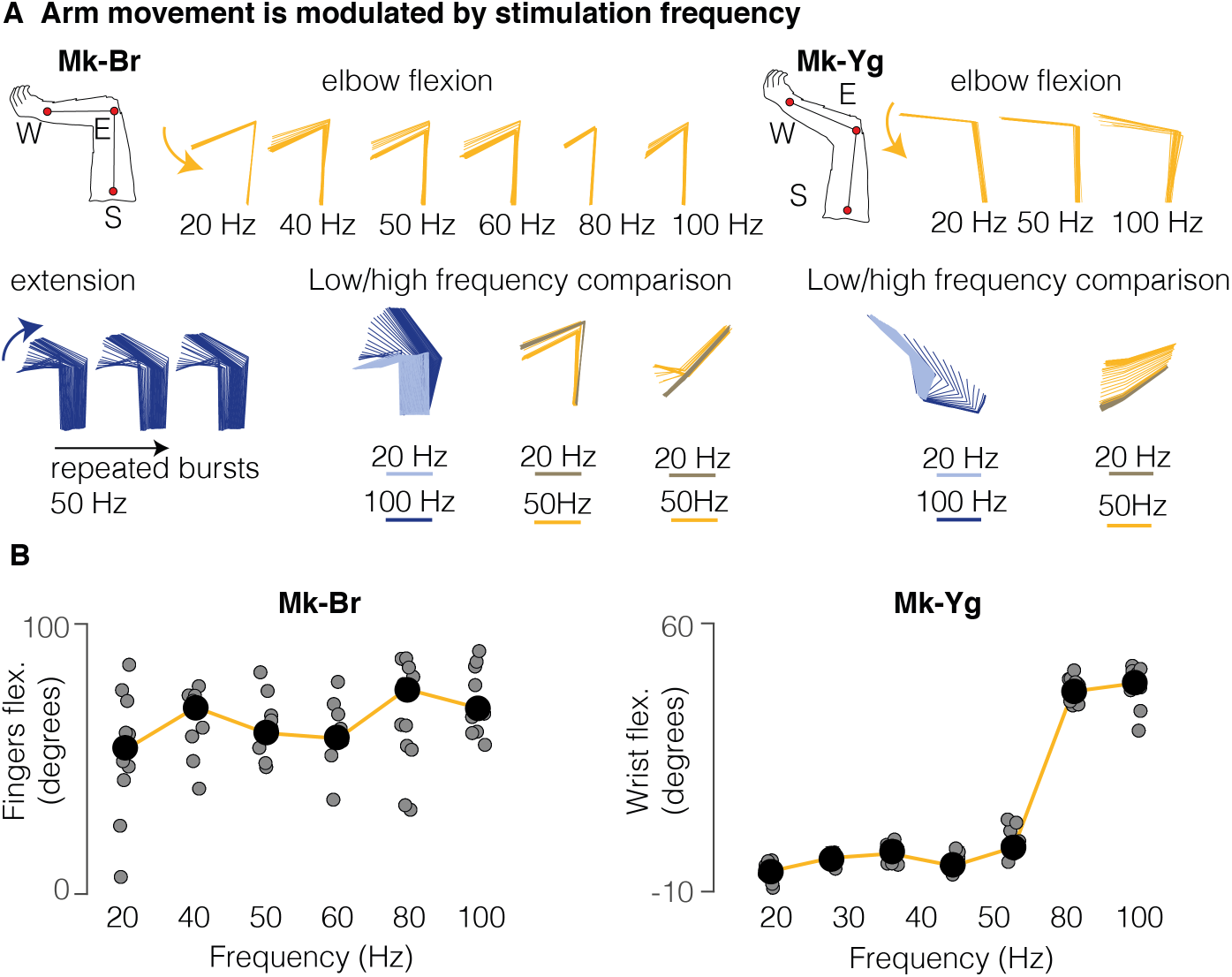
**(A)** Stick diagram schematic of movements elicited by pulse-trains of stimulation in anesthetized conditions. Mk-Br: on the left, arm kinematic obtained by delivering stimulation at different frequencies from contact number 5, on the bottomleft, arm kinematics obtained by repetitive delivery of a burst at 50 Hz; on the bottom right, superimposition of stick diagrams obtained with stimulation at 20 Hz and at higher frequencies (50 or 100 Hz) from different contacts. For Mk-Yg: arm kinematic obtained by delivering stimulation at different frequencies from contact number 2 and superimposition of stick diagrams obtained with stimulation at 20 Hz and at higher frequencies (50 or 100 Hz) from different contacts. **(B)** On the left, finger flexion produced by stimulation at different frequencies from the grasp contact in Mk-Br. Black bullets represent the mean value across different pulse-trains. On the right, wrist flexion obtained by stimulation at different frequencies from the grasp contact in Mk-Yg.

**Extended Data Figure 7.**
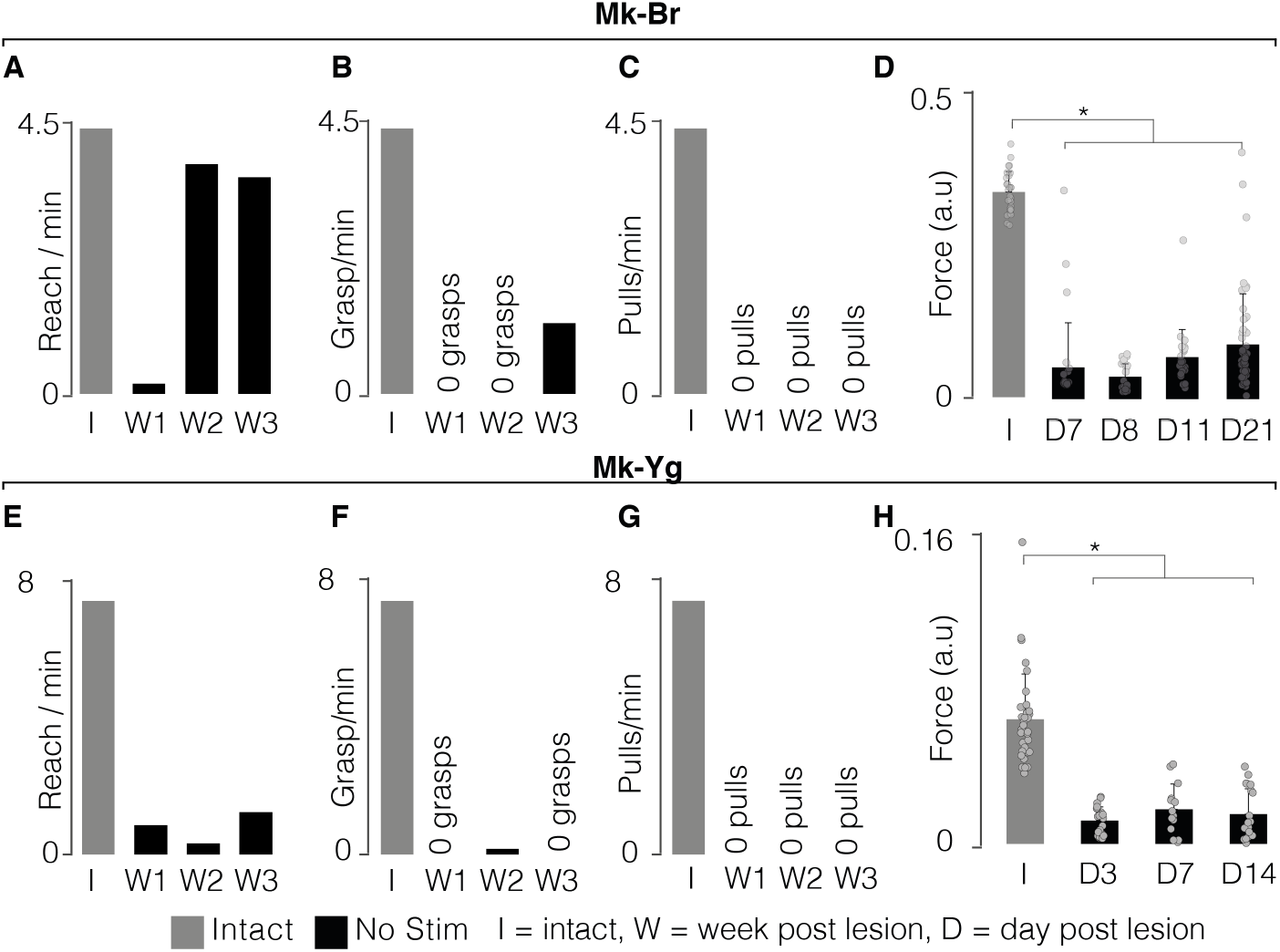
**(A)** Evolution (in weeks) of rates at which Mk-Br performed reach movements after SCI (black), compared to the performances before injury (gray). **(B)** Evolution (in weeks) of rates at which Mk-Br performed grasp movements after SCI (black), compared to the performances before injury (gray). **(C)** Evolution (in weeks) of rates at which Mk-Br performed pull movements after SCI (black), compared to the performances before injury (gray). **(D)** Evolution (in days) of pull force after SCI without stimulation for Mk-Br. Values are plotted as the mean ± SEM. Statistical analysis was carried out with Wilcoxon Ranksum test. **(E)** Evolution (in weeks) of rates at which Mk-Yg performed reach movements after SCI (black), compared to the performances before injury (gray). **(F)** Evolution (in weeks) of rates at which Mk-Yg performed grasp movements after SCI (black), compared to the performances before injury (gray). **(G)** Evolution (in weeks) of rates at which Mk-Yg performed pull movements after SCI (black), compared to the performances before injury (gray). **(H)** Evolution (in days) of pull force after SCI without stimulation for Mk-Yg. Values are plotted as the mean ± SEM. Statistical analysis was carried out with Wilcoxon Ranksum test.

**Extended Data Figure 8.**
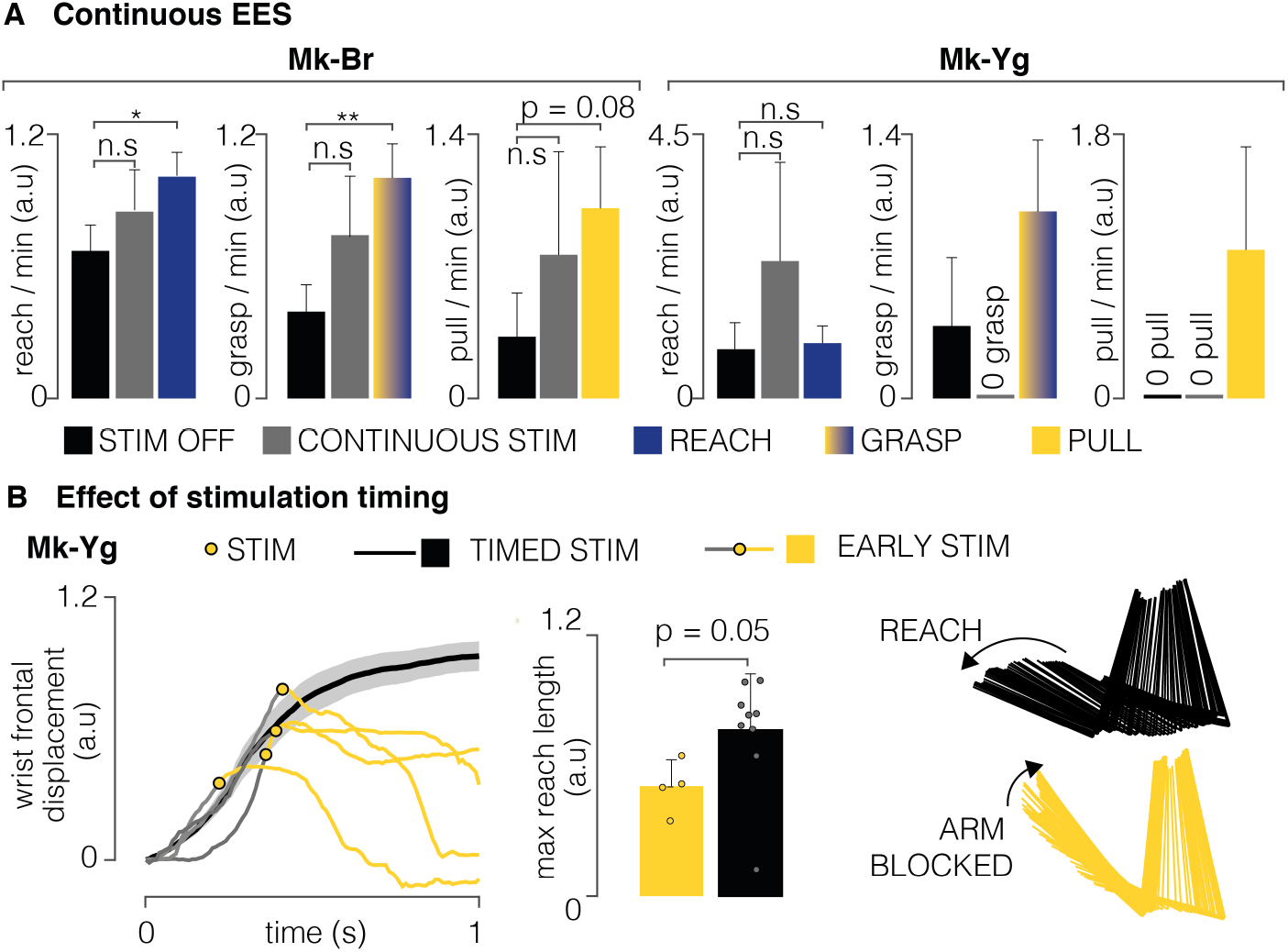
**(A)** Bar plots report the rate of successful movements after SCI, without stimulation (black), with continuous stimulation (gray) and with phase-dependent stimulation (blue or yellow) for Mk-Br and Mk-Yg. Data are presented as mean ± STD and normalized on the mean value in stimulation condition. Statistics was performed with Bootstrap. **(B)** Left: wrist frontal displacement in trials in which pull stimulation was erroneously triggered during reach (gray and yellow), compared to trials in which pull stimulation was not delivered (black). Yellow bullets highlight the instant at which stimulation was delivered: yellow lines highlight the trajectories during and after stimulation. Middle: barplot of the length of the reach movement when pull stimulation was erroneously delivered and when pull stimulation was not delivered. Data are presented as mean ± STD. Right: stick diagram of arm kinematics during reach without (black) and with (yellow) erroneous pull stimulation.

## Materials and Methods

### Animals involved in the study

All procedures were carried out in accordance to the Guide for Care and Use of Laboratory Animals^74^ and the principle of the 3Rs. Protocols were approved by local veterinary authorities of the Canton of Fribourg (veterinary authorization No 2017_04_FR and 2017_04E_FR), including the ethical assessment by the local (cantonal) Survey Committee on Animal Experimentation and final acceptance by the Federal Veterinary Office (BVET, Bern, Switzerland). Three adult female *Macaca Fascicularis* monkeys were involved in the study (Mk-Sa 9 years old, 4.0 kg, Mk-Br 3 years old, 3.4 kg, Mk-Yg 3 years old, 4.0 kg). Animals were not food deprived, could freely access water at any time and were housed in collective rooms designed in accordance to the Swiss guidelines (detention in groups of 2-5 animals in a room of at least 45 m^3^). Rooms were enriched with toys, food puzzles, tree branches and devices to climb and hide, as well as access to an outdoor space of 10-12 m^3^ (see www.unifr.ch/spccr/about/housing). Detailed information on which animals were involved in specific experimental procedures are reported in **Supplementary Table 1.**

### Surgical procedures

For each animal, we performed three surgical procedures, (1) intracortical electrodes implantation, (2) intramuscular electrodes implantation, and (3) epidural implant insertion and spinal cord injury. Mk-Sa deviated from this protocol. Mk-Sa was first implanted with the epidural interface before injury, however an infection occurred and resulted in the explanation of the lead to treat the infection. After recovery, the animal was re-implanted, and lesion performed following the same protocol of Mk-Br and Mk-Yg. All the surgical procedures were performed under full anaesthesia induced with midazolam (0.1 mg/kg, i.m.), methadone (0.2 mg/kg, i.m.), and ketamine (10 mg/kg, i.m.) and maintained under continuous intravenous infusion of propofol (5 ml/kg/h) and fentanyl (0.2-1.7 ml/kg/h) using standard aseptic techniques. A certified neurosurgeon (Dr. Jocelyne Bloch, CHUV, Lausanne, Switzerland) performed all the surgical procedures.

During the first surgical procedure, we implanted multi-microelectrode arrays in the primary motor cortex (M1-42 channels), ventral premotor cortex (PMv-32 channels) and primary somatosensory cortex (S1-42 channels) for a total of 128 channels for Mk-Br and Mk-Yg (Blackrock Microsystems, 400 *μ*m pitch and electrodes tip lengths 1.5 mm 1.5 mm and 1mm for M1, PMv and S1 respectively). Instead, Mk-Sa was implanted with 2 microelectrode arrays of 64 channels each and pitch of 1.5 and 1 mm in M1 and PMd respectively. Functional motor areas of the arm were identified through anatomical landmarks and intra-surgical micro-stimulation. In order to access the brain areas of interest we performed a 20 mm diameter craniotomy and we incised the dura. The arrays implantation was achieved using a pneumatic compressor system (Impactor System, Blackrock Microsystems). A pedestal (*Pedestal A*) was then fixated to a compliant titanium mesh (Medtronic Ti-Mesh) modelled to fit the skull shape and implanted in a previous surgery a few weeks earlier^54^.

During the second surgical procedure we implanted intramuscular electrodes (Teflon-coated stainless-steel wires, Cooner Wire, cat. no. AS631). Mk-Yg received electrodes in the following arm and hand muscles: Deltoid (DEL), Biceps Brachii (BIC), Triceps Brachii (TRI), Extensor Digitorium Communis (EDC), Flexor Carpi Radialis (FCR), Extensor Carpi Radialis (ECR), Flexor Digitorium Superficialis (FDS). Mk-Br received an additional electrode in the Abductor Pollicis Brevis (ABP). Due to practical constraints, Mk-Sa received electrodes only in Biceps Brachii (BIC), Triceps Brachii (TRI) and Flexor Digitorium Superficialis (FDS). In all animals, wires were then connected to an additional pedestal (Pedestal B), fixated to the titanium mesh.

During the third surgical procedure, monkeys were subjected to a lesion at the cervical level (C5/C6) of the spinal cord. The surgeon used a micro-blade to cut approximately one third of the dorsolateral aspect of the spinal cord, in order to interrupt the main component of the corticospinal tract unilaterally. All monkeys retained autonomic functions, as well as limited arm flexion and shoulder adduction capabilities. We monitored the animals for the first hours after surgery and several times daily during the following days. Monitoring scales (score sheets) were used to assess post-operative pain and general health condition during 1-2 weeks. Antibiotics were given immediately after the surgery and then once per day for 10 subsequent days, anti-inflammatory drugs were given once per day for 5 days (Rymadyl 4mg/kg, s.c.; Dexamethasone 0.3mg/kg, s.c.), and analgesic was given twice per day for 5 days (Temgesic 0.01mg/kg, i.m.). Within the same procedure, each monkey received a tailored epidural implant. The implant was inserted in the epidural space of the cervical spinal cord, according to methods described in Schiavone 2020^57^ and Capogrosso 2018^49^. The implant was inserted below the T1 vertebra and pulled until it covered spinal segments from C6 to T1. We performed intra-operative electrophysiology in order to assess and refine the implant positioning so that electrodes are aligned to the animal-specific anatomical features. In particular, we verified that single pulses of stimulation delivered from the most rostral and most caudal electrodes elicited contractions in the BIC and FDS muscles respectively. We re-routed the wires subcutaneously in order to connect them to the *Pedestal B.* All surgical and post-operative care procedures were developed in details in previous reports^49,50^.

### Data acquisition

For Mk-Sa and Mk-Br, we acquired three-dimensional spatial coordinates of arm and hand joints using a 14-camera motion tracking system (Figure 1, Vicon Motion Systems, Oxford, UK) that tracked the Cartesian position of 6 infrared reflective markers (6 to 9 mm in diameter each, Vicon Motion Systems, Oxford, UK) at a 100 Hz framerate. All markers were placed on the left arm, one below the shoulder, three on the elbow (proximal, medial and distal position), and two on the left and right side of the wrist. For each subject, a model of the marker placement was calibrated in Vicon’s Nexus software at the beginning of each experimental session. For Mk-Yg spatial coordinates of arm and hand joints were recorded using two cameras placed parallel to the sagittal and transversal plane of the animal (Vicon Motion Systems, Oxford, UK). The 3D coordinates of the arm and hand joints were extracted using DeepLabCut^75^. Due to the reduced informative content extracted from the camera parallel to the transverse plane, we then only used 2D coordinates on the animals’ sagittal plane. The training set needed for automatic data labeling was created by manually labeling a subset of recorded videos. An investigator was blinded to the experimental condition and was instructed to mark four anatomical landmarks that mirrored the position of markers in Mk-Sa and Mk-Br (shoulder, medial elbow, left and right wrist). Neural signals were acquired with a Neural Signal Processor (Blackrock Microsystems, USA) using the Cereplex-E headstage with a sampling frequency of 30 kHz. Electromyographic signals were acquired with a Behavioral Neurophysiology chronic recording system (RZ2 BioAmp Processor, Tucker-Davis Technologies, USA) at a sampling frequency of 12207 Hz.

### Electrophysiology in sedated monkeys

Monkeys were sedated with a continuous intravenous infusion of propofol (5 ml/kg/h) that minimizes effects on spinal cord stimulation^76^. We delivered single pulses of cathodic, charge balanced, asymmetric square pulses (0.3 ms, 1 Hz) from each electrode contact while recording compound potentials from all implanted arm and hand muscles. Electromyographic signals were acquired with a Behavioral Neurophysiology chronic recording system (RZ2 BioAmp Processor, Tucker-Davis Technologies, USA) at a sampling frequency of 12207 Hz. We then delivered 10 repetitions of pulse trains from each contact, at several frequencies ranging from 20 to 120 Hz. We recorded compound potentials from all implanted arm and hand muscles and arm kinematics through two high resolution cameras (Sony FDR-X3000 Action Cam 4K). Through this procedure we identified three contacts that primarily elicited (1) arm flexors, (2) arm extensors and (3) hand flexors. In a reduced set of trials, we also recorded the force produced by arm flexion through a 10 N range force sensor (Dual-Range Force Sensor, DFS-BTA, Vernier, Beaverton, Oregon, USA). To record the pulling force produced during isometric arm flexion, the hand was fixated to the sensor hook through a string, and the sensor and the elbow were kept in place by two experimenters, in order to optimally capture the strength produced by muscle contraction.

### Behavioral experimental recordings

All animals were trained to perform a three-dimensional robotic reach, grasp and pull task, previously described in detail in (Barra 2019^54^) and briefly recalled here for simplicity. All animals were instructed to wait for a start signal by resting the left hand on a metallic bar. When the “go-cue” was given, monkeys had to reach for and grasp a small spherical object attached to the robot end effector and located in the three-dimensional space. The object was placed approximately 180 mm above the animal seating height, 150 mm far from the shoulder/head coronal plane and 30 mm left of the animal’s left arm. Once animals got a hold on the object, they had to pull it towards their own body until trespassing a virtual spatial threshold. The accomplishment of such virtual threshold was automatically detected by the robot control through online monitoring of the end effector position. Once attained the threshold, monkeys had to let go on the object and go back to the metallic bar. Fruits and vegetables were used to reward successful movements. Animals were trained daily (5 days per week) and every session ended as soon as the animals showed any sign of fatigue or impatience.

For Mk-Sa, data presented in this paper were collected several weeks pre lesion and 1week post lesion, unfortunately a severe infection of the spinal array and EMGs that recurred after day 7 lead to the premature euthanasia of the monkey before the study could be completed, in agreement with the endpoints in our veterinary authorization. For Mk-Br and Mk-Yg data presented in this paper were collected several weeks pre lesion and until 3 weeks post lesion. At the end of week 3 post lesion, Mk-Br had 2 episodes of self-mutilation on the foot ipsi-lateral to the lesion. In consequence we euthanized the animal before the end of the protocol according to the endpoints in our veterinary authorization. As described in the results section, we found post-mortem that Mk-Br had a medial spinal cord contusion at the T3 level. While this lesion did not affect motor control of the legs or the arms, it may have generated neuropathic pain. Mk-Yg could perform the entire protocol without any adverse event, however after day 7, the caudal contact of the spinal interface (E8) identified to promote grasp failed, thus preventing us to perform experiments with optimal stimulation configuration and impacting the efficacy of grasp movements.

### Optimization of EES parameters

To optimized stimulation parameters we exploited the frequency/kinematic relationship that we observed during single contact stimulation (**Figure 3B,E**). We then analyzed single joint movements at different frequencies and contacts and weighted joint excursion angles against movement smoothness^77^, we found that stimulation frequencies of 50-60 Hz (**Extended Data Figure 5**) produced smooth^77^ and full-range movements and maximal forces. Instead, movements elicited at frequencies lower than 40 Hz were too weak to complete a full joint movement while frequencies higher than 60 Hz produced either abrupt movements or incomplete movements (**Extended Data Figure 5A**). Next, we identified among all the tested contacts, those that could consistently elicit arm extension (reach), hand flexion (grasp) and arm flexion (pull) (**Extended Data Figure 5B**). We chose these contacts and 50-60Hz to sustain full arm and hand movement and tested their effect in anesthetized animals by sequentially executing bursts on each of these three contacts. We verified that the sequence triggered whole arm and hand movements that mimicked smooth^77^ and natural multi-joints movements (**Extended Data Figure 5C, Video 1**). Specifically, extension, grasping and pulling movements produced clear EMG bursts as well as robust and smooth kinematics. These stimulation protocols could be triggered by an operator at the beginning of each reach movement or automatically from intra-cortical signals in real-time. Therefore, we verified that movement onset could be detected from intra-cortical signals even after SCI (**Extended Data Figure 5**).

### Stimulation during three-dimensional reach and pull task in injured monkeys

All monkeys were recorded after injury as soon as they could independently move in their housing, feed themselves autonomously and did not show signs of discomfort. This corresponded to 3, 5 and 6 days after injury respectively for Mk-Yg, Mk-Br and Mk-Sa. After injury, the animals were reluctant to perform the task which required intense manual activity by the trainers to encourage them with the use of special positive rewards. Moreover, in consequence of the arm and hand impairments animals were quickly exhausted. As a result, the output of consistent behavior/day was low, and we were able to collect robust data in about 1day/week per animal after SCI. Each session was organized as follows. First, we executed two blocks without stimulation, each of the duration of approximately 2 minutes. During those blocks we visually evaluated the impairment level of the animal and the performance of the brain decoder. Second, we used the brain decoder to trigger specific stimulation patterns. Contacts used to elicit those functions were defined through the experiments described in the previous paragraph and combined together to create stimulation protocols that allowed the animal to perform a full reach, grasp and pull movement.

### Identification and classification of arm movements for kinematic analysis

We defined the movement performed by the animals as composed of three different phases: reach, grasp and pull. The identification of the reach phase was done by marking the moment in which the left hand left the metallic bar to when the hand closed around the object secured to the robot hand effector (the grasp event). The grasp phase was considered to be a window of 100 ms around the moment in which hand closed around the object. The pull phase started from the grasp event and finished when the animal accomplished the task by pulling the object across the virtual spatial threshold and placed the hand back on the resting bar. Events related to the 3 phases of the movement (movement onset: reaching, grasp onset: grasping and release of the object, and pulling) were identified manually by inspecting video recordings from Vicon Motion Systems (Oxford, UK). The same method was applied to mark successful and complete performance of reach, grasp and pull movements as events. A successful reach was defined as a complete extension of the arm that brought the hand at the position of the target (even when grasp could not be performed). A successful grasp was defined as a successful closure of the hand around the target. A successful pull was defined as the accomplishment of a complete flexion movement that brought the target past the virtual spatial threshold. Events were then extracted from Vicon and used to perform analysis on the kinematic of the movements and to train the brain decoder by automatic routines (Matlab 2019b). All the analysis was conducted as blinded experiments.

### Decoding motor states from intracortical signals

We designed a neural decoder that detected reaching and grasping events using intracortical spiking activity. In order to detect spikes, we set a threshold on each channel of −4 times the root-mean-square voltage recorded during a brief period while the monkey was at rest. We estimated firing rates in each of the motor cortical array channels by summing the multiunit spikes with a 150 ms history every 0.5 ms. We used these multiunit firing rate estimates to compute a twenty-dimensional neural manifold capturing the majority of population variance^62^. We projected the spiking activity onto this manifold to calibrate a multiclass regularized linear discriminant analysis decoder^50^ that predicted the labeled timing of reach and grasp events. The decoder used 500 ms of past neural activity and output the probability of observing the reach and grasp events. During calibration, we defined a probability threshold for each event ranging from 0.8 to 0.99 to optimize predictions of the timing of each event using cross-validation. Since the monkeys could not complete the task after SCI, we were unable to consistently acquire labeled training data. We therefore calibrated a decoding algorithm using reaches from a recording session of a healthy monkey. We then manually labeled attempted reaches after SCI by manual inspection of video recordings. Using canonical correlation analysis, we aligned the neural dynamics^78^ preceding reaches on the healthy sessions to the observed neural dynamics preceding attempted reaches after SCI. These aligned dynamics were used to control the decoder trained on the healthy reaches.

We implemented a custom C++ software application running a control suite that used the decoding algorithm to trigger EES stimulation in real-time. The application received neural data over UDP and made predictions using the decoding algorithm at 15 ms intervals. When the output probabilities crossed the defined threshold, the application triggered preprogrammed patterns of EES.

### Analysis of muscle recruitment curves

Electromyographic activity was bandpass filtered between 30 and 800 Hz with an offline 3 ^rd^ order Butterworth filter and stimulus artifact were removed. For each animal, stimulation contact, muscle and stimulation amplitude, we extracted compound potentials from 50ms-long segments of electromyographic activity following a stimulation pulse. We then computed the peak-to-peak amplitude of compound potentials. Since we gave four pulses of stimulation for each selected current amplitude, we averaged across values corresponding to the same stimulation amplitude and represented as the mean recruitment value of each muscle as a function of the injected current. For each muscle, recruitment values have been subsequently normalized by the maximum value obtained for that specific muscle, provided that we obtained response saturation (and therefore maximal contraction) in at least one occasion during the session. In addition, we computed a selectivity index for each muscle^79^.

In order to obtain a comprehensive measure of muscle recruitment for each contact that would allow to compare across animals, we computed, for each animal, each muscle and each contact, an Average Recruitment Index (ARI) as the average of the recruitment values across all stimulation amplitudes used from a specific stimulation site.

To compute muscle recruitment during the delivery of pulse train stimulation, we computed the energy of the EMG signal during the duration of stimulation. We then applied the same normalization procedure described above for single pulse recruitment.

### Analysis of muscle activity during EES

Electromyographic activity was bandpass filtered between 30 and 800 Hz with an offline 3^rd^ order Butterworth filter and stimulus artifact were removed. In all animals we computed the energy EMG signals, for each implanted muscle. Energy of EMG signals during stimulation was computed on each segment in which stimulation was delivered after the animal started a movement attempt, with the formula here below:

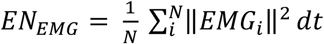

Where *EMG_i_*, is the value of EMG activity at sample *i, N* is the number of samples in the signal and *dt* is the sampling resolution.

Energy of EMG signals without stimulation was computed on each segment in which stimulation was not delivered and the animal started a movement attempt. A movement attempt was defined as an increased EMG activity of the Biceps and Deltoid muscles.

### Analysis of task performance

We computed task performance with two different measures. First, we computed the success rate as the percentage of successful movement across all movement attempts. Successful movements were identified by a blind experimenter as movements performed skillfully and until the end (see above, *Identification and classification of arm movements for kinematic analysis*). Movement attempts were identified as all movements executed in response to a *go* cue and included successful movements too. Second, we computed the task performance frequency as the rate of successful movements per unit of time. In order to do this, we subdivided sessions in time bins of 2 seconds and we marked the presence or absence of successful trials, both with and without stimulation. We then used bootstrap to analyze significance of those results. We normalized all the results by the mean success rate during stimulation.

### Analysis of kinematics performance

We performed Principal Component Analysis on a large set of kinematic features. We computed the features on data segments during the reach phase and the pull phase (see movement identification explained above, section *Identification and classification of arm movements for kinematic analysis*). All kinematic signals were previously low pass filtered at 6 Hz. Segments were not interpolated nor resampled. Before performing PCA analysis, features were centered to have mean 0 and scaled to have standard deviation of 1 (Matlab 2019). The computed features for Mk-Br included: minimum value, maximum value and total excursion of joint angles (shoulder flexion, elbow flexion, and wrist pronation); maximum, minimum and average angular velocity (for the shoulder flexion, elbow flexion and wrist pronation); minimum, maximum and average position along the sagittal, frontal and vertical axis of each arm joint (shoulder, elbow, wrist); maximum minimum and average wrist velocity along the sagittal, frontal and vertical axis; movement smoothness^77^; trajectory length during and time required to complete movements. All the listed features have been computed identically during the reach phase and the pull phase separately and treated as different features. In addition, computed maximal applied three-dimensional pulling force and the average position along the sagittal, frontal and vertical axis of each arm joint (shoulder, elbow, wrist) during grasp.

Since for Mk-Yg we only extracted 2D kinematics on the sagittal plane, the kinematic features for Mk-Yg included: minimum value, maximum value and total excursion of joint angles (shoulder flexion and elbow flexion); maximum and average angular velocity (for the shoulder flexion and elbow flexion); minimum, maximum and average position along the sagittal and vertical axis of each arm joint (shoulder, elbow, wrist); maximum and average wrist velocity along the sagittal and vertical axis; movement smoothness^77^; trajectory length during and time required to complete movements. All the listed features have been computed during the reach phase.

### Processing of cortical signals

We identified spiking events on each channel when the band-pass filtered signal (250 Hz–5kHz) exceeded 3.0–3.5 times its root-mean-square value calculated over a period of 5s. We removed artifacts by deleting all the spikes that synchronously in at least 30 channels. We computed the firing rate of each channel as the number of spikes detected over non-overlapping bins of 10ms. Whenever we showed average firing rate activity, we sorted channels in order of activation in one reference trial, and subsequently applied the same ordering method to all other trials. Finally, we normalized the activity of each channel by its maximum firing rate.

### Comparison of motor cortical activity during EES evoking movement and no movement

To study how motor cortical activity interacted with EES, we analyzed the neural recordings from Mk-Br and Mk-Yg. We identified periods where EES pulse trains produced no discernible movements by setting a threshold on hand velocity. We compared multi-unit neural firing rates on each channel in this period to neural firing rates in the previously identified trials where EES enabled reaching and grasping. First, we counted the number of spikes within the window of stimulation and divided by the duration of stimulation. We then averaged across stimulus repetitions of the movement and no movement conditions and pooled across recording sites in motor cortex.

We next computed instantaneous estimates of multi-unit firing rates on each channel by counting the number of spikes in non-overlapping 20 ms bins and convolving with a gaussian kernel of 50 ms width. We applied Principal Component Analysis (PCA) to compute 10-dimensional neural manifolds spanning this multi-unit population activity^62^. We projected the neural activity onto these manifold axes during the periods where EES evoked either movement or no movement. We then identified periods where the monkey was at rest with no EES, as well as periods where the monkey attempted movements of the arm with no EES. To compare the similarity of neural activity between these conditions, we computed the Mahalananobis distance between activity at rest and the three other periods: EES with movement, EES with no movement, and attempted movements with no EES.

### Histology

Monkeys were deeply anesthetized (lethal dose of pentobarbital, 60mg/kg, injected i.v.) and transcardially perfused with saline (about 200 ml), followed by 3 liters of 4% paraformaldehyde (PFA). Dissected spinal cord were post-fixed in 4% PFA overnight, and then immersed in 30% sucrose solution for 2 weeks. 50*μ*m transverse or horizontal sections were cut using a cryostat and kept in 0.1M PBS azide (0.03%) at 4°C. Primary antibodies were: rabbit anti-Iba1 (1:1000, Wako) and guinea pig anti-NeuN (1:300, Millipore). Fluorescence secondary antibodies were conjugated to: Alexa fluor 647 and Alexa fluor 555 (Life technologies). Sections were coverslipped using Mowiol. Immunofluorescence was imaged digitally using a slide scanner (Olympus VS-120). Lesions were reconstructed using image analysis software (Neurolucida) to trace the lesion over serial sections (200 *μ*m apart).

### Statistical procedures

All data are reported as mean values ± standard error of the mean (s.e.m.) or mean values ± standard deviation (std). The choice is highlighted directly in the figures or in the relative caption. Significance was analyzed using the non-parametric Wilcoxon rank-sum test. In the comparisons shown in Figure 3 we subsequently applied the Bonferroni correction. In only one case (Figure 4A, 4B, 4C), significance was analyzed using bootstrap. The level of significance was set at *p<0.05, **p<0.01, ***p<0.001.

