## Supplementary Data for "Epidural Electrical Stimulation of the Cervical Dorsal Roots Restores Voluntary Upper Limb Control in Paralyzed Monkeys"

| Animal ID | Sex | Age | Weight | SCI | Anesthetized Experiments | Behavioral experiments | Brain array position | Brain - Controlled EES |
|---|---|---|---|---|---|---|---|---|
| **Mk-Sa** | F | 9 Yr | 4 kg | yes | Recruitment curves | Reach only | M1 and PMd | no |
| **Mk-Br** | F | 3 Yr | 3.4 kg | yes | Recruitment curves and single joint movements | Reach and pull | S1, M1 and PMv | yes |
| **Mk-Yg** | F | 3 Yr | 2.9 kg | yes | Recruitment curves and single joint movements | Reach and pull | S1, M1 and PMv | yes |

**Supplementary Table 1**: Identification information, license numbers, characteristics and type of procedure performed for the three monkeys involved in the study

| | **Reach** | | **Grasp** | | **Pull** | |
|---|---|---|---|---|---|---|
| mean ± std (success/min) | No Stim | Stim | No stim | Stim | No Stim | Stim |
| **Mk-Yg** | 0.48±0.13 | 0.53±0.06 | 0.07±0.049 | 0.1±0.03 | 0±0 | 0.015±0.01 |
| **Mk-Br** | 3.08±0.32 | 3.21±0.29 | 0.51±0.13 | 1.06±0.16 | 0.07±0.05 | 0.24±0.07 |
| **Mk-Sa** | 0±0 | 0.13±0.06 | N.A. | N.A. | N.A. | N.A. |

**Supplementary Table 2**: Mean and standard deviation values for non-normalized values of task performance frequency shown in Figure 4C.

**Video 1:** Single-joint movements elicited by pulse trains of EES at different segmental locations. Shoulder abduction: stimulation at C5; Elbow extension: stimulation at C7; Finger flexion: stimulation at T1; reach, grasp and pull sequence: cascade stimulation at C7, T1 and C5.

**Video 2**: Effects of EES on reach movement performance on Mk-Sa. Top left: lateral vision of the animal performing the task; Bottom left: delivered stimulation pulses; Top right: electromyographic activity from Deltoid, Biceps and Triceps muscles; Bottom right: neural activity from M1 and PMd cortex.

**Video 3**: Effects of brain-controlled EES on reach and pull movement performance on Mk-Br Top left: lateral vision of the animal performing the task; Middle left: delivered stimulation pulses; Bottom left: pulling force applied on the robot end effector; Top right: neural activity from S1, M1 and PMd cortex; Bottom right: electromyographic activity from Deltoid, Flexor Carpi radialis and Abductor Pollicis.

**Video 4**: Effects of EES on pull movement performance on Mk-Yg. Top left: lateral vision of the animal performing the task; Middle left: delivered stimulation pulses;

Bottom left: pulling force applied on the robot end effector; Top right: neural activity from S1, M1 and PMd cortex; Bottom right: electromyographic activity from Biceps, Triceps, Extensor Digitorium Communis and Flexor Digitorium Superficialis.

**Video 5**: Effects of a EES burst optimized to recover pull, delivered during a reach movement on Mk-Yg. Lateral view of the animal performing the task.

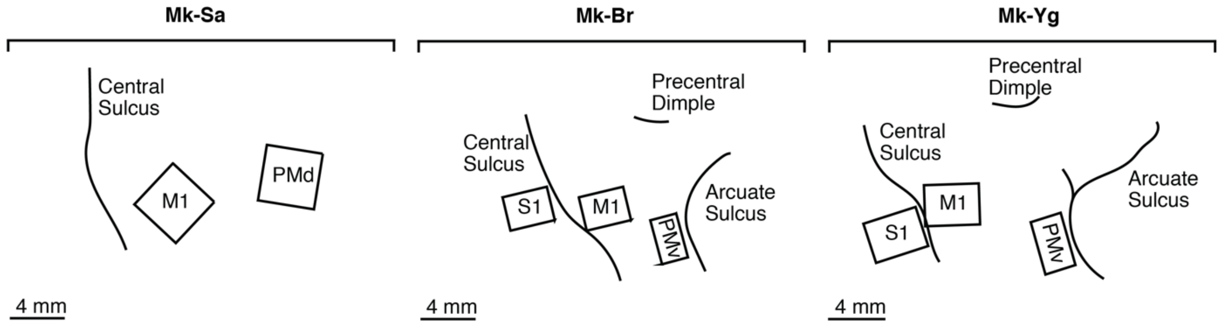


**Supplementary Figure 1: Position of intracortical arrays.** Location of intracortical arrays implanted in the sensory (S1), Motor (M1) and dorsal or ventral premotor (PMd, PMv) cortex in the three monkeys.
